# The motivational control of instrumental performance by nutrient-specific appetites depends on incentive learning

**DOI:** 10.64898/2026.06.14.732213

**Authors:** Douglas J. Roy, Thomas J. Burton, Bernard W. Balleine

## Abstract

Considerable evidence suggests that the motivational control of instrumental action depends on incentive learning; i.e., on the opportunity to learn how the value of the consequences or outcome of an action, (e.g., a specific food) varies under different motivational conditions (e.g., under different degrees of hunger). The current study investigated whether learning the values of high-protein and high-carbohydrate rewards under different degrees of protein and carbohydrate appetite is also necessary for these nutrient-specific appetites to exert control over instrumental performance. Experiment 1 gave differing consummatory experience to whey protein and polycose carbohydrate outcomes under protein and carbohydrate appetite and found that, without the opportunity for incentive learning, the performance of actions earning these outcomes was insensitive to a shift in appetite. However, once the opportunity for incentive learning was provided, the rats increased their instrumental performance on a lever that earned the whey outcome relative to the polycose lever when protein hungry and on the polycose lever relative to the whey lever when carbohydrate hungry. Experiment 2 assessed how these nutrient-specific states exerted this control; whether, once learned, nutrient values were immediately controlled by nutrient appetite or whether this was based on conditional control acquired during experience with the outcomes under different nutrient appetites. We found that exposure to an outcome under a single nutrient-specific state was not sufficient to establish state-specific control. Instead, establishing the conditional control of outcome value required exposure to both the whey and polycose outcomes under both protein and carbohydrate appetites.

## Introduction

There is now considerable evidence that the motivational control of goal-directed instrumental action depends on incentive learning: i.e., on learning how a shift in motivational state affects the value of an action’s goal or outcome (see Dickinson & Balleine, 1994; 2002; Balleine, 2000 for reviews). For example, hungry rats trained to press a lever for a food reward often continue to press when made sated unless they have previously consumed that food when sated (Balleine, 1992). Similarly, incentive learning is required for increases in performance produced by the reverse shift from satiety to hunger (Balleine, 1992; Balleine & Dickinson, 1994) and, indeed, for shifts in the level of thirst (Lopez et al., 1992), between hunger and thirst (Dickinson & Dawson, 1988), in sexual (Everitt & Stacey, 1987; Woodson & Balleine, 2002) and thermoregulatory demands (Hendersen & Graham, 1979), after taste aversion learning (Balleine & Dickinson, 1992) and for drug-induced effects on consumption, e.g., the effect of benzodiazepines on food values (Balleine, Ball & Dickinson 1994).

Nevertheless, while demonstrating the necessity for incentive learning following manipulations of primary motivation, it has long been known that the incentive value of food rewards can also be determined, at least in part, by their nutrient composition. For example, in flavour-nutrient learning, rats’ preferences are biased towards flavours previously associated with nutrients appropriate to their current nutritional state (Sclafani, 1991), including protein (Baker *et al.,* 1987) and lipids and carbohydrates (Davidson *et al*., 1997). The components of protein (amino acids) appear also to relate to specific appetites in ways that affect learning: the behaviour of rats is reportedly influenced by the presence or absence of indispensable amino acids with both foods and cues associated with foods deficient in those amino acids found to be aversive (for a review, see Gietzen, 2021) with similar effects reported in humans (Gibson, Wainwright, and Booth, 1995).

Although these findings appear more relevant to the control of Pavlovian conditioned responses, other evidence suggests that the deprivation of specific nutrients can influence the degree to which those nutrients reinforce instrumental actions. For example, using a progressive ratio procedure, rats have been reported to escalate lever pressing for a high protein pellet to a greater degree when protein-deprived than when non-deprived (Chiacchierini *et al.,* 2022). Similarly, Leatherwood *et al*., (1982; as cited in Ashley, 1985) found that rats trained in a closed economy to press one lever for protein and another for carbohydrate were 10-fold more sensitive to escalation in the response requirement for protein than carbohydrate. However, it has remained unclear whether this evidence of nutrient-specific control of instrumental performance depends on encoding the contingency between the instrumental action and the specific nutrient itself.

We (Roy, Burton, and Balleine, 2026) recently investigated this issue, assessing how nutrient-specific appetites control both the consumption of nutrient-specific rewards and the performance of instrumental actions that earned those rewards. We first gave food-deprived rats the opportunity to learn about the relative values of whey protein and polycose carbohydrate rewards after pre-feeding them a meal high in either protein (i.e., boiled egg or lean steak) or carbohydrate (i.e., cranberries or cupcakes) finding that the rats’ consumption of the whey and polycose solutions was strongly controlled by the pre-feeding: Pre-feeding on steak or egg reduced consumption of the whey vs. polycose, whereas pre-feeding on cake or cranberries reduced consumption of the polycose vs. whey.

We then trained the rats to press two levers, one earning the whey and the other the polycose, before a series of choice tests on the two levers conducted in extinction after the rats were pre-fed either the steak/egg or cranberry/cupcake meals. A similar effect emerged in the instrumental choice extinction tests to that observed in consumption: i.e., after pre-feeding steak or egg, performance on the lever that earned whey was reduced relative to the polycose lever whereas, after pre-feeding cake or cranberries, responding on the lever that earned polycose was reduced relative to the whey lever. Because these effects were observed in extinction, and so without any outcome-related feedback, they are consistent with the claim that the rats encoded the nutrient-specific features of the outcomes associated with the two lever press actions and chose between them based on the value of their consequences; an effect determined by their prior experience with the pre-feeding events. Nevertheless, in the absence of a control group not given the initial consummatory experience, we do not know whether the effects observed on test depended on the opportunity for incentive learning provided by that phase.

The current experiments sought to investigate this issue by establishing: (i) whether incentive learning is required for the control of instrumental performance by nutrient-specific appetites (Experiment 1), and, once established, (ii) whether such control is immediately determined by nutrient appetite or is the product of a state-specific process with conditional control of value acquired via experience of the changing value of specific outcomes under different nutrient appetites (Experiment 2).

### Experiment 1

Experiment 1 used an incentive learning paradigm, similar to that previously reported (e.g., Dickinson & Dawson, 1988; Balleine, 1992) and illustrated in Table 1. Incentive learning was conducted prior to instrumental training when the rats were either protein or carbohydrate hungry, induced by pre-feeding carbohydrate (mixed fruit and maltose) or protein (egg and soy protein), respectively, to food-deprived rats. The instrumental outcomes used during training were whey (protein) and polycose (carbohydrate) solutions. Therefore, prior to instrumental training, the rats were pre-exposed to one or other solution, either the whey (Group Whey-wise) or polycose (Group Sugar-savvy), when protein and when carbohydrate hungry, with these appetites alternating across days. The rats were then trained to press two levers when food deprived, with one lever earning the whey and the other the polycose outcome, after which they were given two choice extinction tests on the two levers, one when protein hungry and the other when carbohydrate hungry. After these tests, a second round of incentive learning was conducted with the rats exposed to the other outcome (e.g., if previously exposed to whey they were now exposed to polycose and vice versa) when protein and carbohydrate hungry, with the appetites again alternated across days, followed by another two choice extinction tests on the levers, one under each appetite.

**Table 1.** Design of Experiment 1. Role of incentive learning in nutrient-specific control

| Group Whey-wise |  |  |  | Instrumental re-training<br>(6 days) |  |
| --- | --- | --- | --- | --- | --- |
| Incentive Learning, Part 1<br>**Whey Exposed**<br>(4 x 2-day blocks) | Instrumental Training<br>(8 days) | Choice test<br>(2 days) | Incentive Learning, Part 2<br>**Polycose Exposed**<br>(4 x 2-day blocks) |  | Choice Test<br>(2 days) |
| **P:** Whey<br>**C:** Whey | **H:** R1→Wh; R2→Pol | **P:** R1 vs R2<br>**C:** R1 vs R2 | **P:** Polycose<br>**C:** Polycose |  | **P:** R1 vs R2<br>**C:** R1 vs R2 |
| Group Sugar-savvy |  |  |  |  |  |
| Incentive Learning, Part 1<br>**Polycose Exposed**<br>(4 x 2-day blocks) | Instrumental Training<br>(8 days) | Choice test<br>(2 days) | Incentive Learning 2<br>**Whey Exposed**<br>(4 x 2-day blocks) |  | Choice Test<br>(2 days) |
| **P:** Polycose<br>**C:** Polycose | **H:** R1→Wh; R2→Pol | **P:** R1 vs R2<br>**C:** R1 vs R2 | **P:** Whey<br>**C:** Whey |  | **P:** R1 vs R2<br>**C:** R1 vs R2 |

**Abbreviations:** **P:** Protein appetite (pre-fed fruit & Maltose); **C:** Carbohydrate appetite (pre-fed egg and soy); **H:** general hunger induced by food-deprivation; **Wh:** Whey protein outcome; **Pol:** polycose carbohydrate outcome; **R1 & R2:** instrumental lever press responses on the left and right lever. *NOTE: the order of presentation of the nutrient events during incentive learning, lever-outcome assignment, and the order of the satiety conditions during the choice tests were fully counterbalanced.*

If incentive learning is required for nutrient-specific appetites to control instrumental performance, then we anticipated that, during Choice Test 1, rats in Group Whey-wise that were exposed first to whey under both appetites would increase performance on the whey lever when protein hungry (pre-fed cabs) relative to when carb-hungry (pre-fed protein). However, because no consummatory experience of the nutrient-specific appetites on the value of the polycose was given to this group, the protein and carb appetites should not modify performance on the polycose lever. Similarly, rats in Group Sugar-savvy, exposed first to polycose under both appetites, should increase responding on the polycose lever when carb-hungry relative to protein hungry whereas nutrient appetite should not control performance on the whey lever. This pattern should change after the second round of incentive learning: i.e., after the Whey-wise group had been exposed to polycose and the Sugar-savvy group exposed to whey when protein and carbohydrate hungry. If incentive learning mediates the effects in the first round of choice tests, then, in the second round of tests, performance on both levers should be controlled by both appetites: i.e., all of the rats should now respond more on the whey than the polycose lever when protein hungry and more on the polycose than the whey lever when carbohydrate hungry. Of course, these predicted effects should not emerge if our previously observed effects of nutrient appetite on instrumental performance were direct and so not a function of incentive learning.

## Methods

### Subjects

Subjects were 16 male and 16 female outbred Long Evans rats, 250-350 g prior to start of experiment, obtained from Animal Services at the University of New South Wales (Randwick, NSW). For all experiments, rats were housed in opaque plastic boxes in groups of 4 (males and females in separate boxes) in a climate-controlled colony room and maintained on a 12 h light/dark cycle (lights on between 07:00 and 19:00). All experimental stages occurred during the light portion of the day. Water and standard lab chow were continuously available prior to the start of the experiment. All experimental procedures were approved by the Animal Ethics Committee at the University of New South Wales and are in accordance with the guidelines set out by the American Psychological Association for the treatment of animals in research.

### Food deprivation

Three days before the start of behavioural procedures, the rats were handled, weighed daily, and placed on a food deprivation schedule. This consisted of giving rats free access to water bottles in the home cage but no lab chow. The macro-nutritional profile of the lab chow was 18.1% crude protein, 65.2% carbohydrate, and 5.1% fat, with the protein being primarily from casein, and carbohydrates derived from corn and cellulose. An hour after each training or test session, rats were given combined access to lab chow and water for 1 hour in their home cages before the deprivation conditions were reimposed. Their weights were monitored daily to ensure each remained above 85% of their pre-procedure body weight throughout.

### Apparatus

For all instrumental procedures, the apparatus consisted of 16 operant chambers, each enclosed in a sound- and light-attenuating shell (MED Associates, Vermont USA). Each chamber was equipped with two pumps that were fitted with syringes and delivered either 0.2 mL of whey protein or polycose solution into separate wells within a recessed magazine, and a pellet dispenser that also delivered 45 mg grain food pellets (Bioserve) into the magazine. Retractable levers were used for instrumental training, with rats presented with one at a time during training sessions while the other was retracted, and with both presented simultaneously during choice tests sessions. Levers were mounted on the front panel, one on the left and the other on the right of the magazine. A fan located on the back wall of the shell provided constant background noise (∼60 dB). The chambers were equipped with a sound card that delivered 90 dB 3000 Hz clicks and 90dB white noise. The walls were clear Plexiglas, and the floor was a stainless-steel grid, below which was an aluminium tray for collecting subjects’ droppings. Computers running MED Associates proprietary software (Med-PC) controlled all experimental events and recorded lever presses and magazine entries.

### Nutrient-specific events

The outcomes used as rewards in Experiment 1 were whey protein, mixed by adding 30 g of whey protein concentrate to 100 mL water plus 1 mL of vanilla essence for flavouring (yielding ∼93 kcal per 100 grams of the outcome), and polycose (maltodextrin), mixed by adding 30 g of polycose to 100 mL water plus 1 mL of vanilla essence (∼88 kcal per 100 grams of the outcome). These substances and concentrations were chosen because of comparability in their metabolic effects (see Roy et al., 2026). The high protein meals consisted of fried egg, prepared by mixing equal parts whole egg with egg whites to raise the protein content relative to non-protein energy (fat) and create a uniform distribution of nutrients, presented in ceramic ramekins and a 10% solution of unflavoured soy protein isolate powder mixed in water. These were presented in the home cage in a ramekin and drinking bottle respectively for 30 min. Soy protein isolate was chosen here because, like whey, it is a high-quality protein with similar amino acid profiles while differing in sensory properties. The satiety induced by soy protein should therefore translate into reduced value of the whey protein reward despite differing markedly in sensory characteristics. The high carbohydrate meals consisted of a medley of dried diced fruits (including banana, cranberries, sultanas, peaches, pears, apple, and apricot) presented in the same kind of ramekin as the egg, plus a maltose solution mixed with water to a concentration of 10% and presented in drinking bottles similar to those used for the soy protein mentioned above. Pre-feeding and incentive learning sessions were conducted in the rats’ home cages to ensure the familiarity of the pre-feeding environment. The average quantity consumed during these pre-feeding events was calculated by dividing the total amount consumed by the number of subjects per cage.

### Incentive learning Part 1

Prior to the start of this phase of the experiment, the rats were divided into two groups: Whey-wise (n=16; 8 male and 8 female) and Sugar-savvy (n=16; 8 male and 8 female). Rats in the Whey-wise group were given alternating sessions in which they could consume the whey outcome from a bottle in the home cage immediately after either the high protein meal (egg+soy) or after the high carbohydrate meal (fruit+maltose). These experiences were given on alternate days over a total of eight days; i.e., four of each type of pre-feeding (see Table 1). Exactly the same experience was given to the Sugar-savvy group except that polycose was given after both pre-feedings instead of whey. Total consumption during the pre-feeding and incentive learning (meals and drinks) were measured before and after each session and the difference divided by the number of rats to establish average consumption.

### Instrumental training

All rats were food-deprived throughout instrumental training. To ensure similar acquisition rates in each group we first trained the rats to press each lever separately with both levers delivering grain pellets. Accordingly, rats were first given 2 days of magazine training with 20 grain pellets delivered on a random-time 30 sec schedule, after which they were trained across 8 consecutive days of instrumental pre-training with pellets delivered first on a continuous reinforcement schedule (days 1 to 4) and then on a random ratio 5 schedule on days (5 to 8).

Rats were then trained to lever press for the whey and polycose outcomes with one outcome delivered on one lever (either left or right) and the other on the alternate lever (right or left, counterbalanced). Levers were trained one at a time so that, for example, if the right lever was presented and could be pressed to earn polycose in one session, then in another session the left lever was presented to earn the whey protein outcome. Each session terminated after rats earned a total of 20 outcomes or after 20 minutes had elapsed. Rats were trained to press for whey protein and polycose on alternating days (order counterbalanced), such that they only had a 20-minute session training them to press one lever for whey one day, a different lever for polycose the next, and so on, until rats had received 8 training sessions with each lever. Each training session was separated by a day so that rats did not confuse or misattribute metabolic effects across types of reward.

Rats were initially trained on a continuous reinforcement schedule. When an individual earned all 20 outcomes of both type for two consecutive sessions, they were escalated to a reward schedule of random-ratio 5 (RR-5) again until all outcomes were earned in two consecutive sessions, and then to RR-10 for the remainder of training.

### Choice test 1

After instrumental training, two outcome devaluation choice tests were conducted. Half of the rats in each group were pre-fed with the egg+soy protein with the remainder pre-fed the fruit+maltose meal in the home cage. Based on studies assessing protein and carbohydrate metabolism (Chiacchierini et al., 2025; Grove et al., 2022) pre-feeding consisted of continuous free access to one of the two meals for 30 minutes in the feeding chambers. After pre-feeding, the rats were immediately placed in the operant chambers, both levers were extended, and they were free to press both levers in a choice extinction test for 5 minutes, meaning that the levers could be pressed but no outcomes would be delivered. This was important because the test sought to assess whether rats were choosing to work for an outcome based on both an expectancy of the value of that outcome given current appetite conditions, and the knowledge of which specific reward each action was expected to produce. The next day, rats in both groups were given a second choice extinction test after exposure to the other nutrient meal; i.e., if they were given the first test after consuming the protein meal then they were given the carbohydrate meal and vice versa, with the choice test conducted in the same manner as described above.

### Predictions

If these tests revealed that instrumental performance for nutrient-specific rewards is mediated by incentive learning then we predicted that: the Whey-wise group should press the whey lever more than the polycose lever when protein hungry (i.e., when pre-fed carbohydrate) whereas the Sugar-savvy group should press the polycose lever more than the whey lever when carbohydrate hungry (i.e., when pre-fed protein). In contrast, because incentive learning under the differential nutrient appetites had not been provided for polycose in Group Whey-wise, or whey in Group Sugar-savvy, then choice performance shouldn’t differ in these groups when carbohydrate or protein hungry, respectively.

### Incentive learning Part 2 after a change in outcome

To provide the groups with full information on the value of whey and polycose under the protein and carbohydrate appetites, we next gave all of the rats a second set of incentive learning experiences but now on the alternate outcome: i.e., Group Whey-wise were now exposed to polycose both after pre-feeding egg+soy, in one session, and after pre-feeding fruit+maltose, in the alternating session, whereas Group Sugar-savvy were now exposed to whey both after pre-feeding egg+soy, in one session, and after pre-feeding fruit+maltose, in the alternating session across 4, 2-day blocks.

### Instrumental re-training and testing 2

After incentive learning, all rats were given 6-days of instrumental re-training with the two lever press actions earning the same outcomes as in initial training delivered on the RR-10 schedule. After this training the rats were given two choice extinction tests exactly as described above with choice between the two levers assessed after pre-feeding protein or carbohydrate in alternation (counterbalanced).

### Predictions

If the motivational control of instrumental performance by nutrient-specific appetites depends on incentive learning then the predictions for the two groups should be the same for the second round of testing after the second round of incentive learning: i.e., now that all of the rats have been exposed to both outcomes in both nutrient states, we predict a full cross over-interaction: rats in *both* groups should choose the whey over the polycose lever when protein hungry (i.e., after being pre-fed fruit+maltose) and the polycose lever over the whey lever when carbohydrate hungry (i.e., after being pre-fed egg+soy).

### Statistical analyses

A primary set of three hypotheses were directly tested by conducting statistical analyses of the devaluation choice test data using either 2-way or 3-way ANOVAs (some of which were mixed-models) followed by planned comparisons. Although the specifics of each test are stated in the corresponding results section, the ANOVAs typically included factors of Appetite (protein vs. carbohydrate), Outcome (whey vs. polycose) and Instrumental Incentive Learning (IIL; before vs. after full IIL).

Firstly, with groups collapsed, we hypothesised that the extent of Instrumental Incentive Learning (IIL) would interact with the Nutrient-Driven Devaluation (NDD) effect (i.e. the interaction between Appetite and Outcome), which would be revealed by a significant 3-way interaction between the factors of Appetite, Outcome and IIL. Next, decomposing the data by Group and IIL, we predicted that prior to full IIL, only the reward with which rats had received sufficient IIL experience with should be successfully devalued by sating rats on the relevant appetite i.e., presses on the protein lever should be suppressed by protein pre-feeds compared to carbohydrate pre-feeds in Group Whey-wise (but not Group Sugar-savvy, and vice versa). This would manifest as significant Appetite-Outcome interactions. Finally, following full IIL, presses on both type of lever should be appropriately sensitive to pre-feeds regardless of group assignment i.e. the full Nutrient-Driven Devaluation effect (NDD). This would also be evidenced by a significant Appetite-Outcome interaction.

## Results and Discussion

### Incentive learning

#### Group Whey-wise

Consumption and instrumental training data provide important context for interpreting the data in the choice tests. Figure 1 shows consumption of whey in Group Whey-wise across exposure sessions after each type of pre-feeding in the first (Figure 1a) and second (Figure 1b) incentive learning episodes. There was a clear elevation in consumption of whey after fruit+maltose vs. egg+soy pre-feeding that increased across sessions during the first incentive learning episode and a similarly clear elevation in consumption of polycose after egg+soy vs fruit+maltose pre-feeding across sessions during the second incentive learning episode. These data were analysed using three-way ANOVA comparing consumption by Outcome (whey vs. polycose), Appetite (protein vs. carbohydrate), and Session finding: no main effects of Outcome (*F*(1, 3) = .378, *p* = .582, 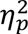 = .112), Appetite (*F*(1, 3) = .555, *p* = .510, 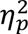 = .156), or Session (*F*(1, 3) = 1.118, *p* = .368, 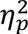 = .272) but a significant Outcome × Appetite interaction *F*(1, 3) = 83.669, *p* = .003, 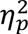 = .965 and three-way Outcome × Appetite × Session interaction *F*(1, 3) = 21.628, *p* = .019, 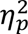 = .878, suggesting that the two way interaction emerged from repeated experience. Other interactions between Session × Outcome (*F*(1, 3) = 4.349, *p* = .128, 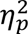 = .592) and Session × Appetite (*F*(1, 3) = .882, *p* = .417, 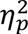 = .227) were not significant.

**Figure 1.**
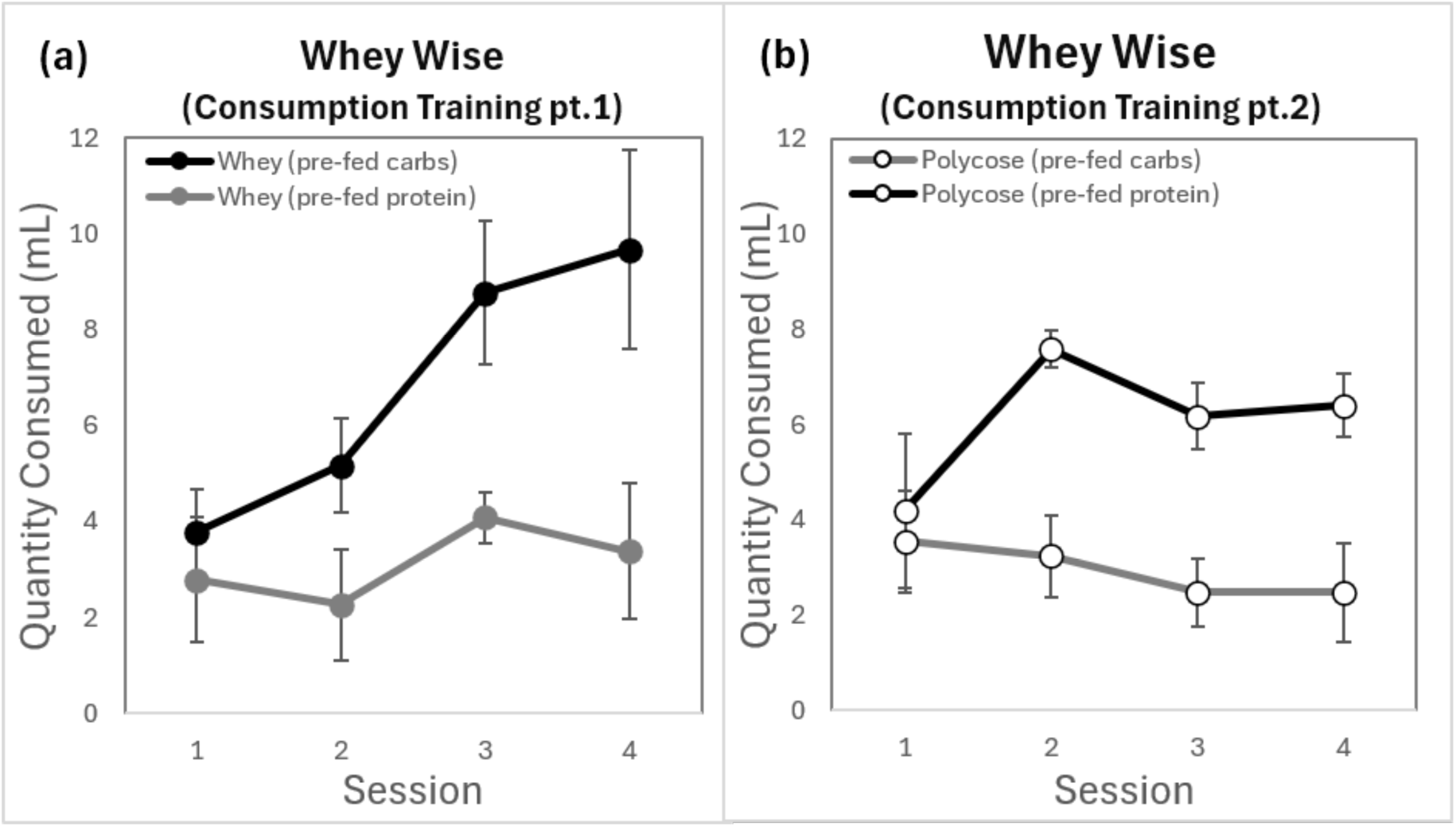
Incentive learning in Group Whey-wise. Average volume of **(a)** whey and **(b)** polycose consumed after protein vs. carbohydrate pre-feeding during incentive learning episodes 1 and 2. (Error bars = ± 1 SEM)

#### Group Sugar-savvy

Similar effects emerged in Group Sugar-savvy across incentive learning episodes, shown in Figures 2a and 2b. Again, a three-way ANOVA revealed no main effects of Outcome (*F*(1, 3) = 1.702, *p* = .282, 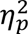 = .362), Appetite (*F*(1, 3) = 3.734, *p* = .149, 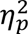 = .555), or Session (*F*(1, 3) = 6.501, *p* = .084, 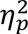 = .684), but found a significant Outcome × Appetite interaction, *F*(1, 3) = 99.103, *p* = .002, 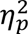 = .971, that again interacted with Session; i.e., the three-way interaction: *F*(1, 3) = 33.504, *p* = .010, 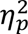 = .918. The interactions Session × Outcome (*F*(1, 3) = 1.357, *p* = .328, 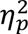 = .312) and Session × Appetite (*F*(1, 3) = 4.715, *p* = .118, 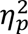 = .611) were again not significant. Therefore, the consumption patterns during both incentive learning episodes in both groups were consistent with the conclusion that experience under the alternating appetites was sufficient to produce incentive learning regarding the value of each outcome under the specific appetites.

**Figure 2.**
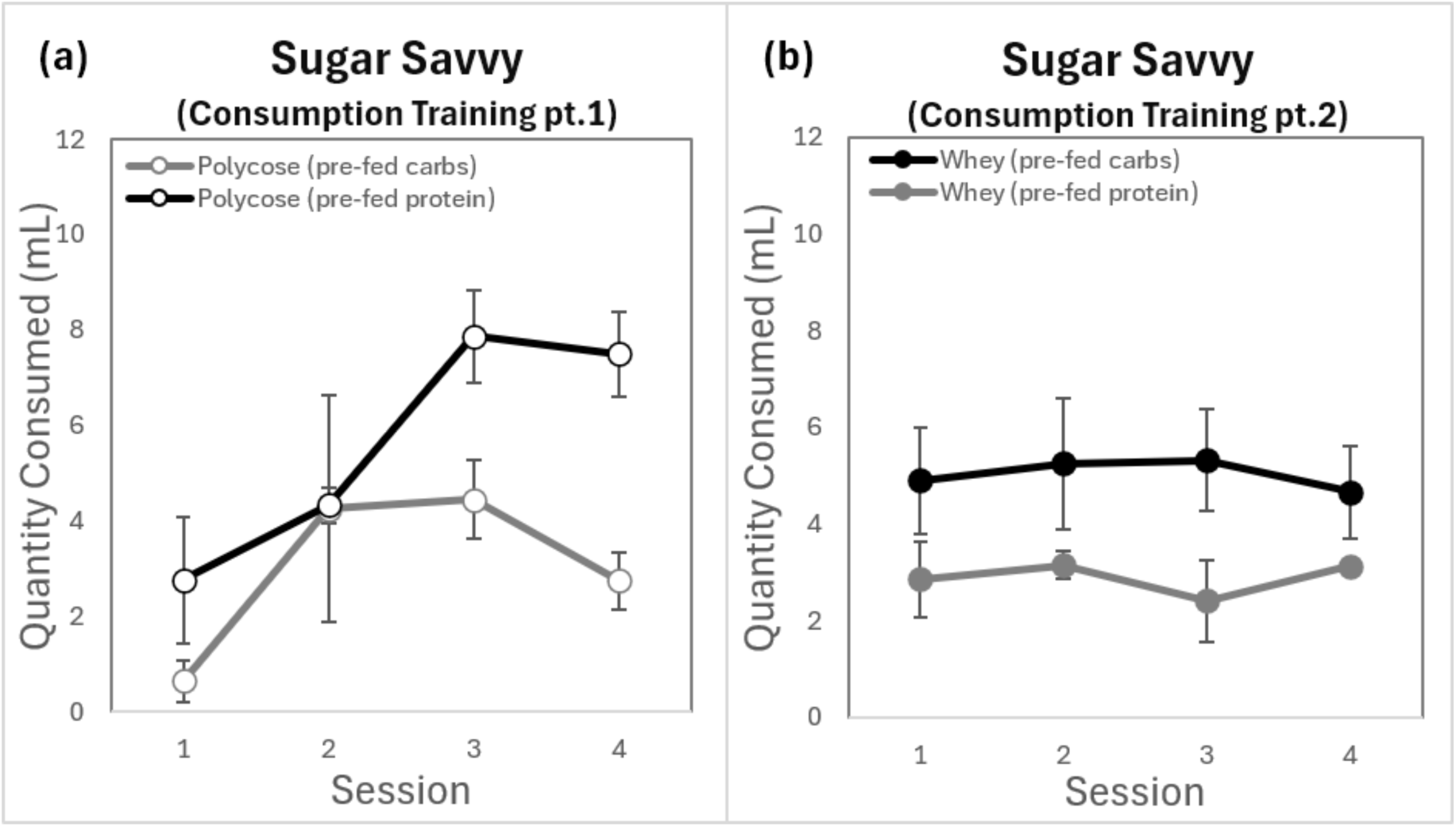
Incentive learning in Group Sugar-savvy. Average volume of **(a)** whey and **(b)** polycose consumed after protein vs. carbohydrate pre-feeding during incentive learning episodes 1 and 2. (Error bars = ± 1 SEM)

### Instrumental training

Instrumental training data for each group before and after the two episodes of incentive learning are presented for Groups Whey-wise and Sugar-savvy in Figures 3 and 4, respectively. As is clear from these figures, rats in Group Whey-wise pressed slightly but consistently more for polycose in both training periods, although this slight bias was not evident in Group Sugar-savvy. To check whether baseline responding before each test was significantly biased across incentive type and experimental conditions, instrumental acquisition data were analysed with a mixed-model ANOVA, with two levels of Group (Whey-wise vs. Sugar -Savvy) two levels of Outcome (whey protein vs. polycose), two levels of IIL training Phase (before vs. after full IIL), and two levels of Session (lever press rates averaged during the first two and during the last two sessions to compare early versus late for each training phase). This confirmed successful instrumental training across both cohorts. There was no significant main effect of Outcome (*F*(1, 30) = .340, *p* = .564, 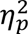 = .011) or Group (*F*(1, 30) = .942, *p* = .340, 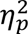 = .030), and no significant interaction between Outcome and Group, (*F*(1, 30) =3.964, *p* = .056, 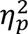 = .117). A main effect of session block confirmed increased performance across training blocks, *F*(1, 30) = 188.60, *p* <.001, 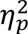 = .863. This was qualified by a significant Phase x Session interaction, presumably reflecting savings from prior instrumental training *F*(1, 30) = 7.206, *p* = .012, 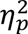 = .194. That is, while rats in the initial training phase demonstrated a steep increase from a low baseline to asymptotic levels, re-training performance commenced at a high level immediately following the first round of testing, resulting in distinct acquisition trajectories across phases despite reaching similar final performance peaks. No other interactions reached statistical significance (all p’s > .129).

**Figure 3.**
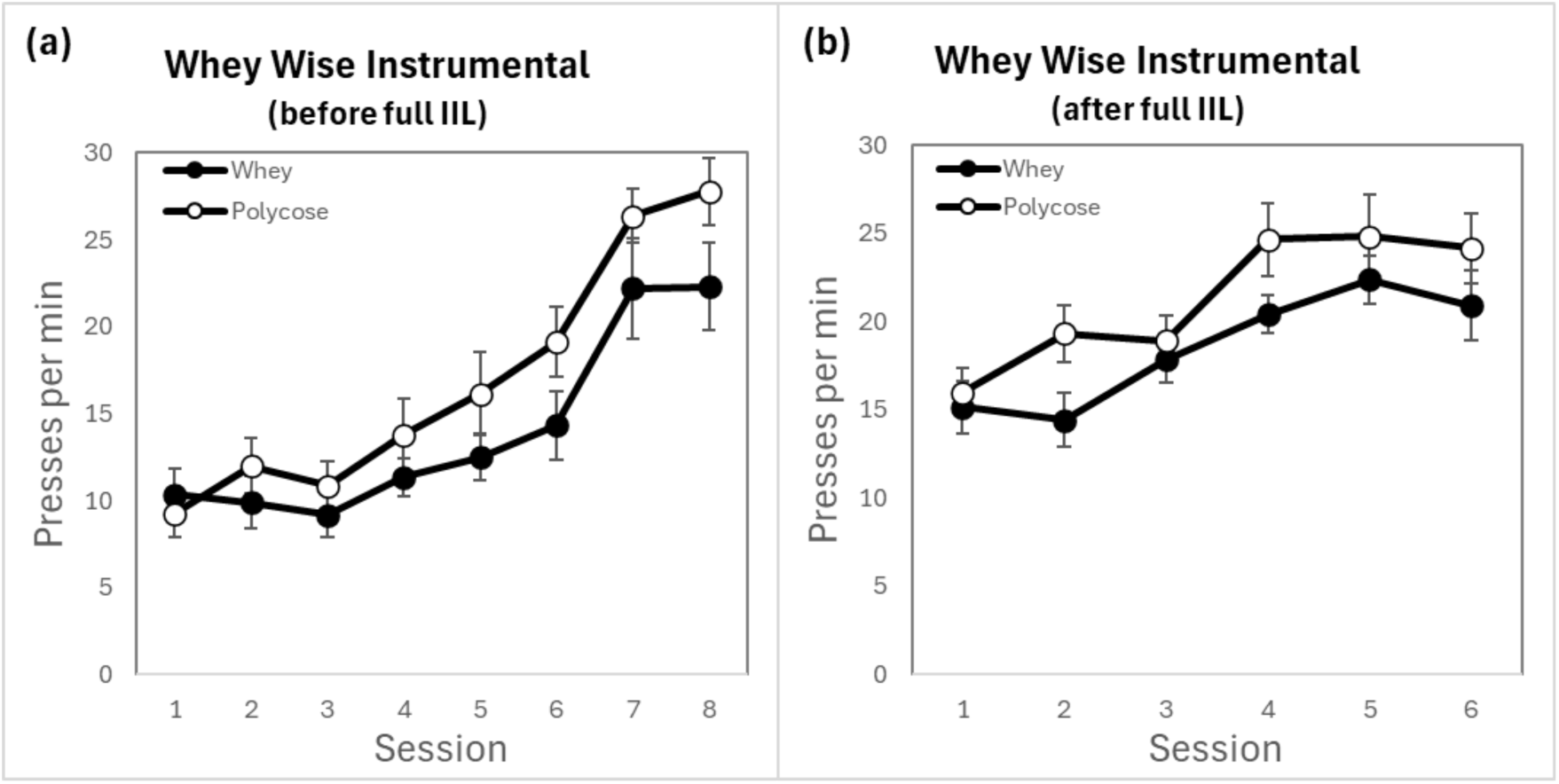
Responses on the whey and polycose levers during the instrumental training and re-training periods for Group Whey-wise. Average lever press rates for each session during acquisition training are shown for **(a)** the first period of training before full incentive learning and the first set of choice tests and **(b)** re-training after full incentive learning and before the second set of tests. (Error bars = ±1 SEM).

**Figure 4.**
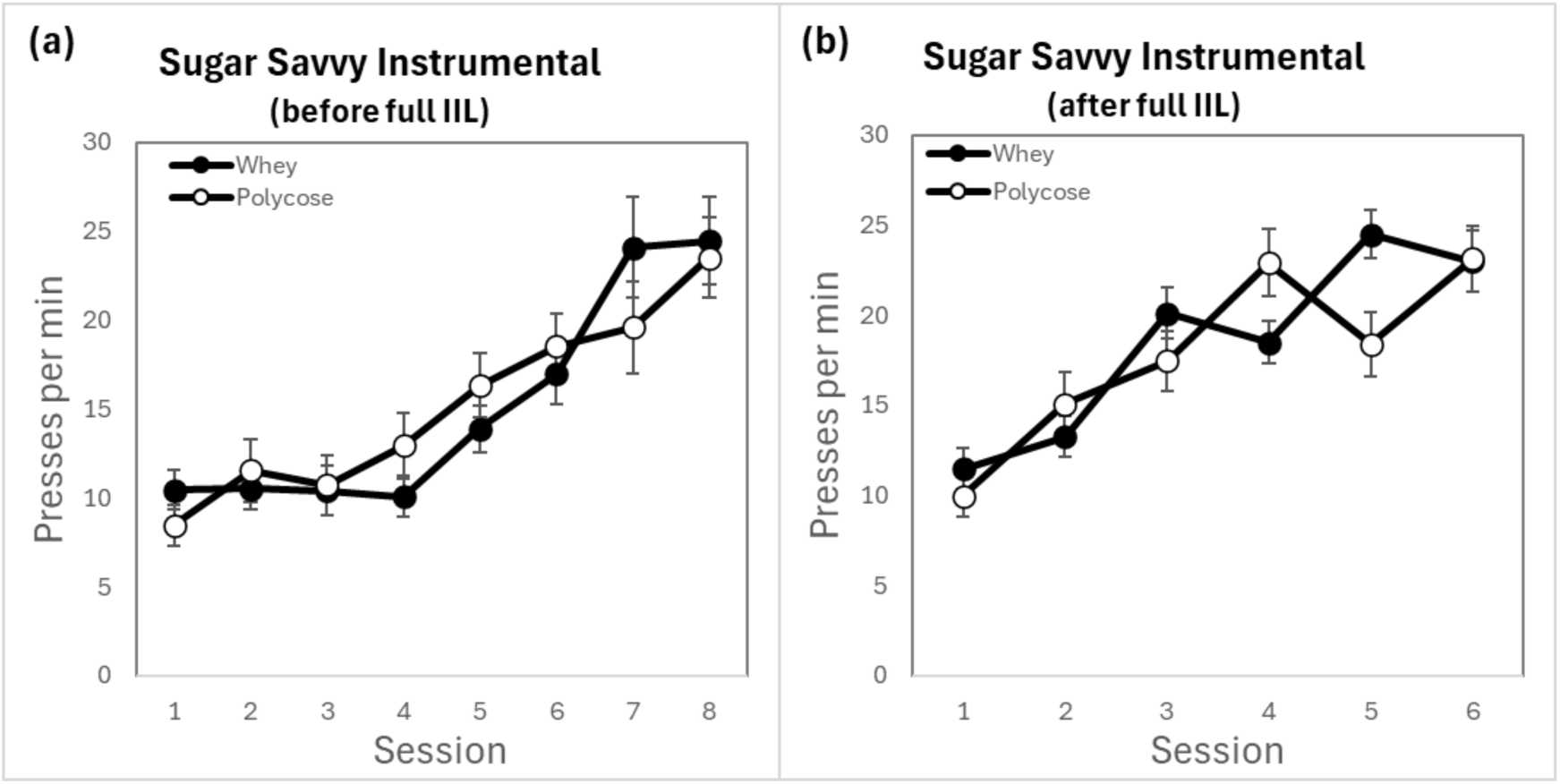
Responses on the whey and polycose levers during the instrumental training and re-training periods for Group Sugar-savvy. Average lever press rates for each session during acquisition training are shown for **(a)** the first period of training before full incentive learning and the first set of choice tests and **(b)** re-training after full incentive learning and before the second set of tests. (Error bars = ±1 SEM).

### Choice tests conducted after partial and full incentive learning

The critical data from the choice tests are presented first for each group separately and for each of the two sets of choice tests; i.e., after partial and full incentive learning experience. We then present the data collapsed across the group factor to reveal the general role of incentive learning in the control of instrumental performance by nutrient-specific appetite.

#### Group Whey-wise after partial incentive learning

Performance in the first set of choice tests after partial incentive learning in Group Whey-wise is shown in Figure 5. The control by nutrient appetite appeared clearest in the test conducted protein hungry, with a mild increase early in the test on the whey lever vs. polycose lever that was not evident in the test conducted carbohydrate hungry. Indeed, there was a clear failure to observe any direct control of performance by carbohydrate appetite with pressing on the polycose lever being greater under the protein than the carbohydrate appetite. This pattern suggests that responding on the whey lever under the protein appetite was largely a product of incentive learning.

**Figure 5.**
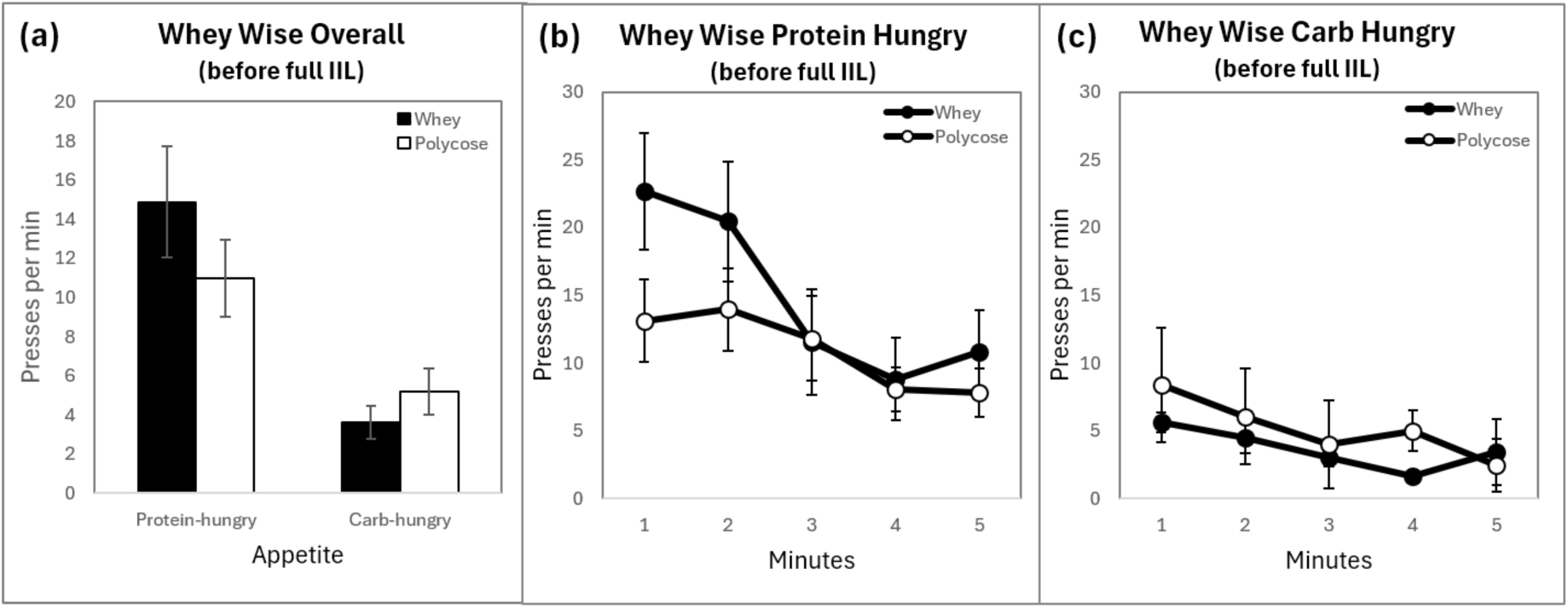
Lever press rates on the whey vs. polycose levers during the extinction choice tests in Group Whey-wise after partial incentive learning. **(a)** Mean lever press rates for the whey protein lever vs. polycose lever across the two appetite conditions. Average press rates for each consecutive minute of the test when **(b)** protein hungry, and **(c)** carbohydrate hungry (error bars = ±1 SEM).

The statistical analysis supported this description. A two-way ANOVA was conducted on mean rates of lever pressing with factors of Outcome, separating the levers by the outcome earned during training (whey vs. polycose), and Appetite, separating responding during the test protein hungry (i.e., after pre-feeding with fruit+maltose) from that carbohydrate hungry (i.e., after pre-feeding egg+soy). This analysis revealed no main effect of Outcome (*F* (1, 15) =.312, *p* =.585, 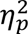 = .020) but a significant main effect of Appetite, *F* (1, 15) =22.501, *p* <.001, 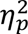 = .600) and, more importantly, a significant Outcome x Appetite interaction *F* (1, 15) =7.484, *p* =.015, 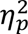 = .333. Simple effects analyses conducted on the interaction revealed a significant effect of Appetite on performance on the whey lever, *F* (1, 15) =22.25, *p* <.001, 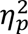 = .597, with rats responding at significantly higher rates under protein than carbohydrate appetite. It also revealed a significant effect of Appetite on the polycose lever, however, here responding was not congruent with appetite: responses on the polycose lever were also higher under the protein than the carbohydrate appetite, *F* (1, 15) =12.239, *p* =.003, 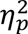 = .449. Nevertheless, there was no effect of Outcome, either in the test conducted protein hungry, (*F* (1, 15) =1.170, *p* =.296, 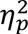 = .072), or carbohydrate hungry (*F* (1, 15) =1.708, *p* =.211, 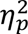 = .102).

#### Group Whey-wise after full incentive learning

The data from the second set of choice extinction tests conducted after full incentive learning are presented in Figure 6. Although incentive learning appears to have had only a small influence when exposure was only given to whey, the overall pattern became far clearer after full incentive learning. After exposure to the polycose in the different nutrient appetites the rats now appeared to press more on the polycose lever when carbohydrate hungry than when protein hungry. Similarly, the bias in responding on the whey vs. the polycose lever under the protein appetite was also clearer. The statistics supported this description. As previously, the average lever press rates during the choice extinction tests after full incentive learning for Group Whey-Wise were analysed by a two-way ANOVA with factors of Outcome and Appetite. Although neither the main effects of Outcome (*F* (1, 15) =.078, *p* =.784, 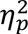 = .005) or Appetite (*F* (1, 15) =3.755, *p* =.072, 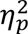 = .200) were significant, there was a significant Outcome x Appetite interaction *F* (1, 15) =13.374, *p* =.002, 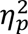 = .471. Simple effects analyses found that rats responded more on the whey lever in the protein hungry test than in the carb-hungry test, *F* (1, 15) =11.022, *p* =.005, 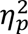 = .424, and more on the polycose lever in the carb-hungry test than the protein hungry test, *F* (1, 15) =7.026, *p* =.018, 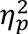 = .319. However, although the Whey-wise group responded more on the polycose than the whey lever in the carb-hungry test, *F* (1, 15) = 15.802, *p* =.001, 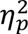 = .513, responding on the two levers in the protein-hungry test did not differ significantly, (*F* (1, 15) =3.065, *p* =.100, 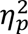 = .170). Generally, therefore, the pattern of result in the choice tests for the Group Whey-wise suggests that control by the nutritive appetites emerged selectively after partial incentive learning, being limited to the whey lever when protein hungry, but emerged more completely when the opportunity for incentive learning was given for both outcomes under both nutritive appetites.

**Figure 6.**
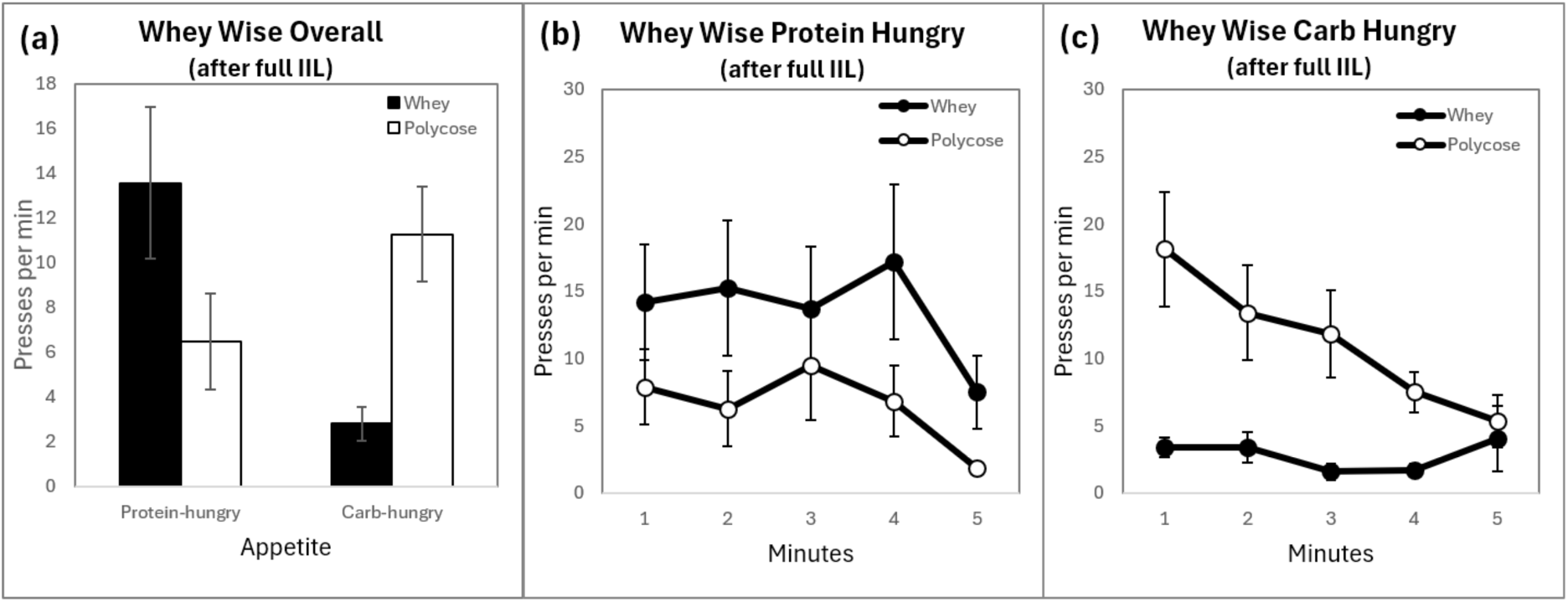
Lever press rates on the whey vs. polycose levers during the choice extinction tests in Group Whey-wise after full incentive learning. **(a)** Mean press rates for the whey protein lever vs. polycose lever across the two appetite conditions. Average press rates for each consecutive minute of the test when **(b)** protein hungry, and **(c)** carbohydrate hungry (error bars = ±1 SEM).

#### Group Sugar-savvy after partial incentive learning

Generally speaking, the pattern of results observed in Group Whey-wise was also observed in Group Sugar-savvy. After partial incentive learning, control by nutritive state was limited to the test conducted carbohydrate hungry, summarised in Figure 7. A two-way ANOVA conducted on the overall lever press rates during the choice extinction test found no main effect of Outcome (*F* (1, 15) =1.631, *p* =.221, 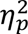 = .098), or Appetite (F (1, 15) =1.736, *p* =.207, 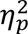 = .104) but revealed a significant Outcome x Appetite interaction, *F* (1, 15) =11.649, *p* =.004, 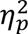 = .437. Simple effects analyses revealed that, whereas press rates on the whey lever were not significantly affected by appetite (*F* (1, 15) =1.436, *p* =.249, 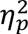 = .087), those on the polycose lever were greater when carbohydrate hungry than when protein hungry, *F* (1, 15) =8.163, *p* =.012, 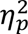 = .352. Furthermore, in the carbohydrate hungry test, rats pressed more on the polycose than the whey lever, *F* (1, 15) = 8.781, *p* =.010, 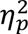 = .369, whereas this difference was not significant in the protein hungry test (*F* (1, 15) =1.213, *p* =.288, 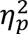 = .075).

**Figure 7.**
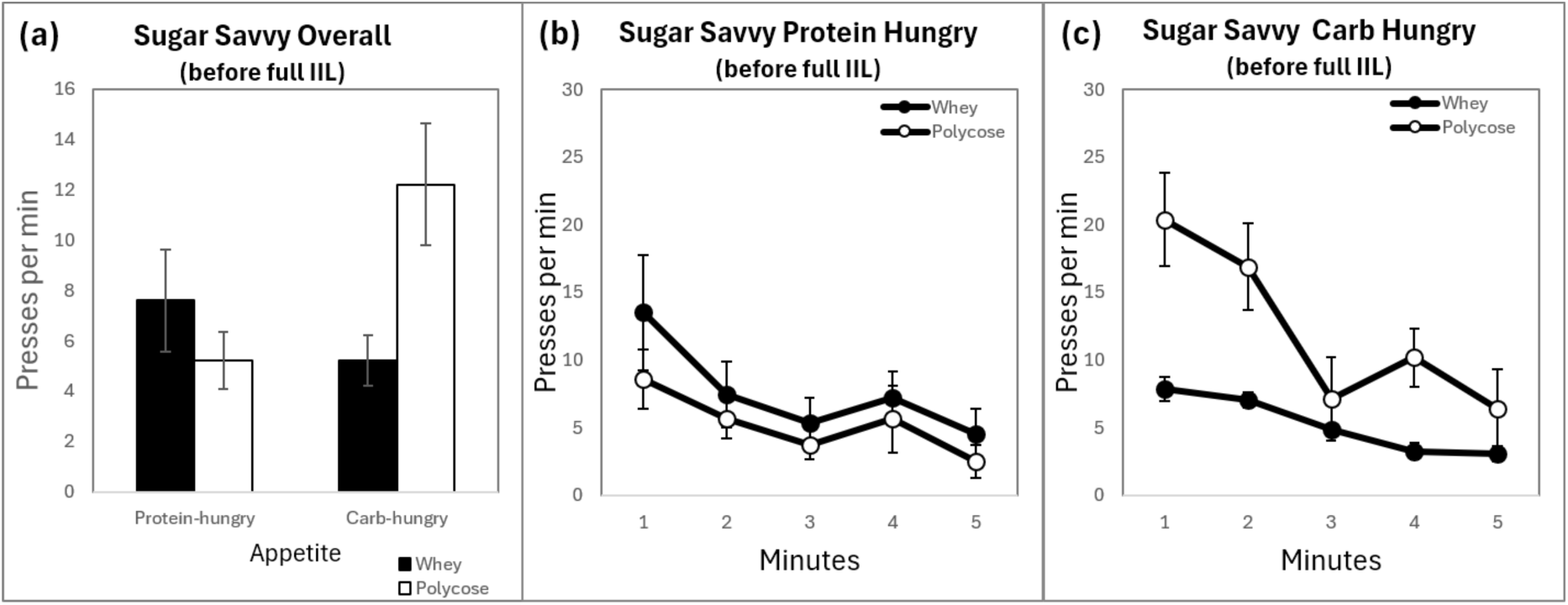
Lever press rates on the whey vs. polycose levers during the extinction choice tests in Group Sugar-savvy after partial incentive learning. **(a)** Mean press rates for the whey protein lever vs. polycose lever across the two appetite conditions. Average press rates for each consecutive minute of the test when **(b)** protein hungry, and **(c)** carbohydrate hungry (error bars = ±1 SEM).

#### Group Sugar-savvy after full incentive learning

The results of the test conducted after exposure had been given to both the whey and polycose outcomes under the protein and carbohydrate hunger states is presented in Figure 8. These results were very similar to those observed in Group Whey-wise (Figure 6). and indeed, were statistically similar too. The two-way ANOVA found no main effect of Outcome (*F* (1, 15) =.683, *p* =.421, 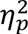= .044) *or* Appetite (*F* (1, 15) =3.929, *p* =.066, 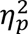 = .208) but revealed a significant Outcome x Appetite interaction, *F* (1, 15) =16.296, *p* =.001, 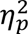 = .521. Simple effects revealed that the rate of pressing on the whey lever was greater when protein hungry than when carb-hungry, *F* (1, 15) =13.232, *p* =.002, 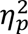 = .469, whereas the rate of pressing on the polycose lever was greater when carbohydrate hungry than when protein hungry, *F* (1, 15) = 8.300, *p* =.011, 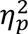 = .356. During the protein hungry test rats pressed the whey lever more than the polycose lever, *F* (1, 15) = 21.924, *p* <.001, 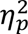 = .594. whereas, during the carb-hungry test they pressed the polycose lever more than the whey lever (*F* (1, 15) = 4.754, *p* =.046, 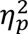 = .241).

**Figure 8.**
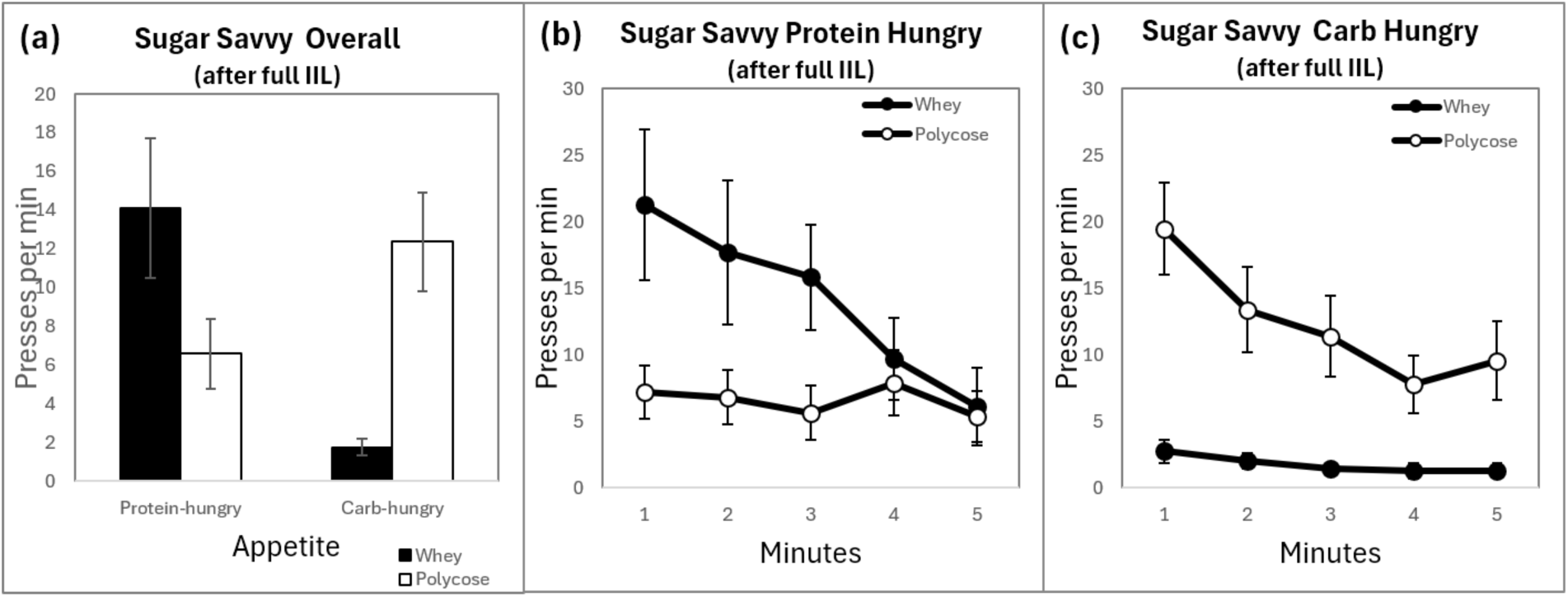
Lever press rates on the whey vs. polycose levers during extinction choice tests in Group Sugar-savvy after full incentive learning. **(a)** Mean lever press rates for the whey protein lever vs. polycose lever across the two appetite conditions. Average press rates for each consecutive minute of the test when **(b)** protein hungry, and **(c)** carbohydrate hungry (error bars = ±1 SEM).

#### Results of outcome devaluation tests before and after incentive learning (overall)

When the above findings for Whey Wise and Sugar Savvy rats are considered together, it is compelling to conclude that IIL was required for rats to adjudicate choice between actions based on the nutritive value of reward according to appetite. They demonstrate that rats can differ in how much they know about two types of rewards at the same time, being “savvy” from IIL about one while remaining “naïve” about the other, and that this can change with subsequent experience. Interestingly, when rats knew about the relation between the whey protein reward and appetite but not the polycose, they seem to have generalized and treated the polycose like it was a source of protein (but the opposite result did not occur in the group with the opposite set of experiences).

#### The combined effect of shifts in nutrient appetite before and after incentive learning

Finally, we combined the data of the two groups across the tests based on whether an action previously delivered an outcome congruent with current nutritive state on test (i.e., a valued outcome) or incongruent with that state (i.e., a devalued outcome), and whether the opportunity for incentive learning had been provided through exposure to the outcome in the congruent state prior to the test (Savvy) or not (Naive). Performance of the rats during the choice extinction tests conducted after partial incentive learning is shown in Figure 9a, which can be compared with the results when the rats were re-tested after full incentive learning with both rewards in Figure 9b. Press rates clearly differed on the valued vs. the devalued lever when rats had been given appropriate incentive learning (Savvy) but did not differ without this exposure (Naive). In contrast, when incentive learning had been given to both outcomes, rats now showed a marked consistency, with performance now determined solely by the value of the outcome under the nutritive appetites. To reveal this most clearly, we combined performance on the levers in both groups during the final choice extinction tests to reveal the motivational control of instrumental performance by nutrient-specific appetites after incentive learning had been given - Figure 9c. There it is most clearly revealed that, after full incentive learning had been given, responding on the whey and polycose levers during the extinction tests depended on the nutrient-specific appetite in which the rats were tested.

**Figure 9.**
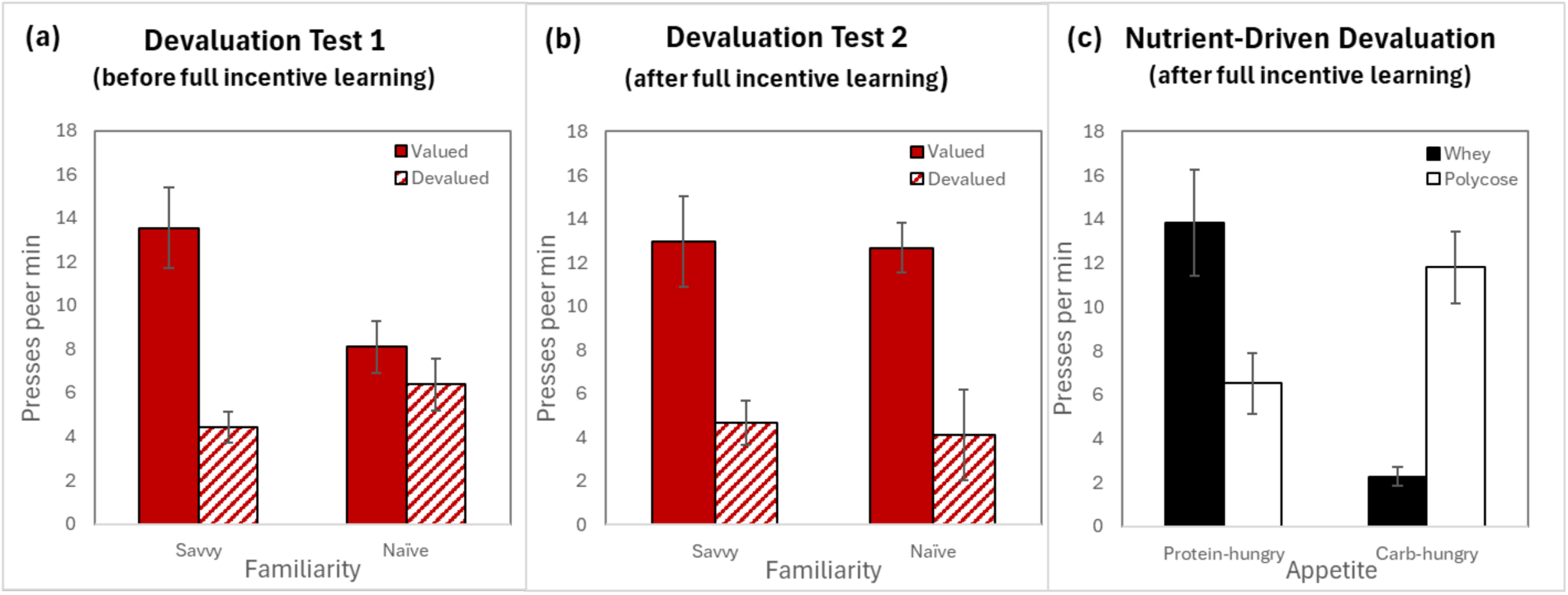
Overall lever press rates averaged across the Whey-wise and Sugar-savvy groups during the extinction choice tests. Mean lever press rates on the valued (congruent with nutritive state) and devalued (incongruent with nutritive state) levers when the outcome had been devalued after **(a)** partial and **(b)** full incentive learning. **(c)** Mean press rates by lever identity and appetite conditions after full incentive learning for both groups combined (error bars = ±1 SEM).

We analysed the data presented in Figures 9a and 9b together in a single three-way ANOVA using factors of Value (valued vs. devalued), Incentive Learning (Savvy vs. Naive) and Test (Test 1 vs Test 2) and critically found a significant three-way Value x Incentive Learning x Test interaction, *F* (1, 31) = 7.015, *p* =.013, 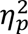 = .190. This appeared to depend on different degrees of incentive assessed during the tests: whereas we found a Value x Incentive learning interaction in Test 1 *F*(1, 31) = 15.975, *p* < .001, 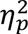 = .340, no such interaction emerged in Test 2, (*F*(1, 31) = .014, *p* = .905, 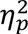 = .000).

Finally, we analysed the data, presented in Figure 9c, using a three-way ANOVA incorporating a factor of Group (Whey-Wise vs. Sugar-savvy), with factors of Lever (whey vs. polycose) and Appetite (protein hungry vs. carb-hungry). This analysis found that there was no significant main effect of Outcome (*F* (1, 30) =.524, *p* =.475, 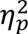 = .017) or Group (*F* (1, 30) = .007, *p* =.936, 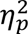 = .000), nor did Group feature in any significant interactions (all *p*’s > .678). There was a significant effect of Appetite, with rats pressing on average more when hungry for protein (*M* = 10.18, *SD* = 8.36) than when hungry carbohydrate (*M* = 7.05, *SD* = 5.14)*, F* (1, 30) = 7.679, *p* =.010, 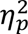 = .204. There was a significant Outcome x Appetite interaction *F* (1, 30) =29.656, *p* <.001, 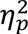 = .497. Simple effects analyses found that rats responded more on the whey lever in the protein hungry test (*M* = 13.84, *SD* = 13.74) than in the carb-hungry test (*M* = 2.28, *SD* = 2.49), *F* (1, 30) = 24.954, *p* <.001, 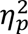 = .446, and more on the polycose lever in the carb-hungry test (*M* = 11.82, *SD* = 9.21) than the protein hungry test (*M* = 6.52, *SD* = 7.81), *F* (1, 30) =15.763, *p* <.001, 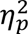 = .337. Rats also responded more on the polycose than the whey lever in the carb-hungry test, *F* (1, 30) = 30.00, *p* <.001, 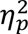 = .552, and less to the whey lever than the polycose lever during the protein-hungry test, *F* (1, 30) = 7.794, *p* =.009, 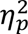 = .201.

Generally, therefore, Experiment 1 found that nutrient-specific appetites can exert an exquisite level of control over the performance of instrumental actions that earn commodities relevant to those appetites but only after incentive learning has been given, i.e., only after those commodities have been consumed under the nutrient-specific appetites, allowing their values under the specific appetites had been learned. Importantly, we found no evidence of any implicit or direct control of instrumental performance by nutrient-specific appetite in the absence of incentive learning, and this was true both of protein and carbohydrate appetites. Therefore, this experiment supports the claim that the motivational control of instrumental performance by nutrient-specific appetites depend on incentive learning.

### Experiment 2

Prior analyses of incentive learning have suggested that associating an outcome with an emotional response is sufficient to establish its current incentive value with the prevailing motivational conditions determining the intensity and valence of that response (Balleine, 2001). However, in addition to establishing the value of the outcome, these analyses also point to the role of motivational state in the control of that value (Dickinson & Balleine, 1994). Thus, when the motivational conditions change, the experienced change in the value of the outcome generates the need to bring that new value under the conditional control of the current motivational state both to avoid ambiguous value encoding and to ensure the appropriate value is applied given the prevailing motivational conditions. For example, when a particular food is experienced hungry, rats assign a high value to it. However, they tend to persist in applying that high value even when they become sated – the so-called resistance to satiation effect (Morgan, 1974; Podelsnik & Shahan, 2009) – unless they experience the new value in that sated state. It is this experience that allows them to learn that the different values are conditional on motivational state: a high value is assigned when hungry and a low value when sated. As a consequence, the tendency to generalise these values across states is reduced, ensuring the appropriate value is assigned to the food based on their current state.

Although our results are consistent with those of previous assessments of motivational control, we do not currently know if this type of incentive learning applies to nutrient-specific appetites. This is because, in Experiment 1, we gave the rats exposure to each of the outcomes under a protein and a carbohydrate appetite (in alternation) during incentive learning, and then in a generally food-deprived state during instrumental training. As a consequence, the rats had considerable experience with the value of at least one of the two outcomes from the very start of the experiment and, therefore, had the opportunity for those states to exert conditional control over its incentive value. In Experiment 2, we sought to evaluate if exposure to the outcome in more than one state is necessary for nutrient-specific appetites to exert control over instrumental performance.

To achieve this, in Experiment 2 we employed similar methods to those used in Experiment 1, except that we exposed the rats to the two outcomes in only one motivational state during incentive learning. Using the design presented in Table 2, we trained the rats to press two levers, one for whey and the other for polycose as in Experiment 1. However, unlike Experiment 1 we trained the rats in either a protein hungry or a carbohydrate hungry state from the very start of the experiment. In this way we hoped to ensure that the values of the two outcomes remained constant: one group, Whey-high, were trained protein hungry and so the value of the whey protein was high and the polycose carbohydrate was low, whereas, the other group, Whey-low, was trained carbohydrate hungry and so the value of whey protein was low and the polycose carbohydrate was high. We then assessed whether these values were controlled by motivational state by giving the rats two choice tests, one when protein hungry and the other when carbohydrate hungry. Based on prior findings, we anticipated that, without the opportunity for nutrient-specific appetites to establish conditional control over the values of the whey and polycose outcomes, the rats would apply the values as they were experienced during training regardless of their current motivational state. If, however, the values of the whey and polycose outcomes are immediately and directly determined by nutrient-specific state then both groups should respond at a higher rate on the whey lever than the polycose lever when protein hungry and at a higher rate on the polycose lever than the whey lever when carbohydrate hungry.

**Table 2.**
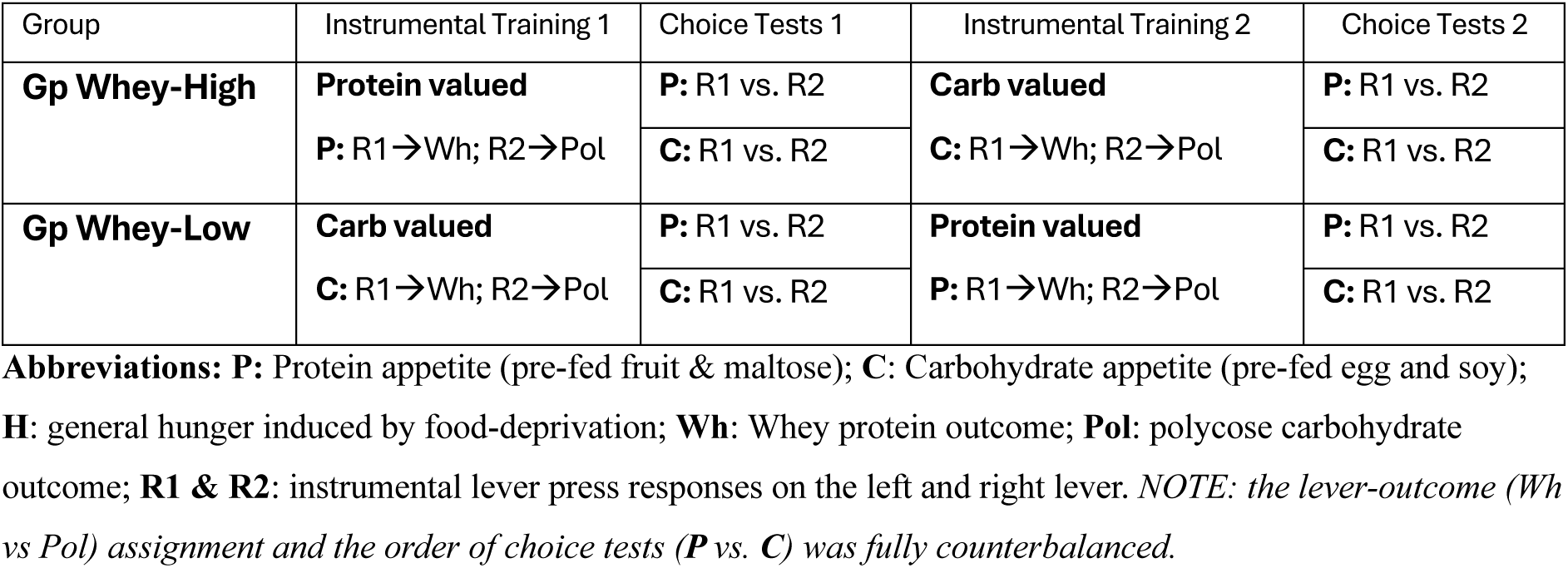
Design of Experiment 2. Conditional control of nutrient-specific values

The results from the first test suggested that conditional control was not immediately established and so we next sought to induce such control by giving the rats the opportunity to consume the outcomes after a shift in motivational state: i.e., the rats continued to earn the two outcomes on the two levers as before but were now shifted to the alternate state, with the Whey-high group now trained under a carbohydrate appetite, thereby reducing the value of whey and increasing the value of polycose, and the Whey-low group trained under a protein appetite, reducing the value of polycose and increasing the value of whey. We then gave the rats a second set of tests under the two nutrient-specific appetites to establish whether this subsequent training experience was sufficient to bring the values of the whey and polycose outcomes under the control of the nutrient-specific state. If this were the case then we predicted that the rats in both groups would now respond at a higher rate on the whey lever than the polycose lever, when protein hungry, and at a higher rate on the polycose lever than the whey lever, when carbohydrate hungry. Of course, no such pattern of results should emerge if this training experience was not sufficient to induce such conditional control.

## Methods

### Subjects

Subjects were 8 male and 8 female outbred Long Evans rats, 250-350 g prior to start of experiment, obtained from Animal Services at the University of New South Wales (Randwick, NSW) and housed as described in Experiment 1. Three days before the start of behavioural procedures the rats were weighed and placed on a food deprivation schedule on which they were given free access to water but only sufficient chow each day to maintain them at 85% of their pre-procedure body weight. Rats were assigned to one of two experimental groups: (1) Group Whey Low (*n*=8; 4 male and 4 female), and (2) Group Whey High, (*n*=8; 4 male and 4 female), with reference to the incentive value of the whey protein reward during the initial 10-day training phase.

### Apparatus

For all instrumental procedures, the apparatus used in Experiment 2 were the same as those described in Experiment 1.

### Nutrient-specific events

The same nutrient-specific events were used in Experiment 2 to those described in Experiment 1. Prior to the beginning of instrumental procedures, rats were given two opportunities to sample both types of meal to overcome any neophobic reactions. The whey and polycose solutions that served as the instrumental outcomes were mixed and flavoured as described in Experiment 1.

### Procedure

#### Instrumental training

The design of the experiment is presented in Table 2. Rats were given 2 days of magazine training and 10 consecutive days of instrumental pre-training with presses on both levers rewarded with grain pellets (two sessions per day, each with either the left or right lever, order counterbalanced across days). They were initially trained on a continuous reinforcement schedule (days 1 to 4) but escalated to a RR-5 reinforcement schedule (days 5-to 10) from the 5^th^ day of training to facilitate subsequent learning that actions produced specific outcomes under conditions where their motivation to work for rewards might be diminished by the pre-feeding manipulations.

Rats were then divided into two groups and trained to press one lever for whey protein and another for polycose carbohydrate delivered on a RR-5 schedule. Each instrumental training session ran for 20 minutes or until 10 rewards were earned. The rats were then immediately given a training session with the other lever and outcome (session order counterbalanced between rats and across days). If rats earned all their outcomes from both levers two days in a row, they were escalated to an RR-10 reinforcement schedule. For this instrumental phase (lasting ten days), rats were initially trained under motivational conditions in which the appetite for protein versus carbohydrate was manipulated by pre-feeding. One reward was always high in value while the other was always low throughout training. One group of rats, Whey-high (*n=*8), were trained when protein hungry, which was achieved by feeding them the fruit followed by maltose, whereas the other group, Whey-low (*N=*8), was trained only when they were hungry for carbohydrate, achieved by pre-feeding them the egg and soy protein meal.

#### Choice extinction tests

For the test sessions, the rats were first given access to either a high protein or a high carbohydrate meal, followed initially by a 5-minute test conducted in extinction and then a 10-minute test in which both rewards could be earned on a RR-5 schedule. The next day, the test was repeated but following the type of meal they had not received on the first test; i.e., if the rats were protein sated on test day 1, then they were carbohydrate sated on test day 2 and vice versa.

#### Retraining and testing

After the initial period of training and testing, the rats were given four days of retraining but under the opposite motivational conditions in place during initial training: i.e., rats in Group Whey-high were now trained under a carbohydrate appetite whereas those in Group Whey-low were now trained under a protein appetite. This reversal of training conditions was intended to provide the animals with the opportunity for incentive learning after the change in state and to encourage them to bring the various values of the whey and polycose outcomes under the control of the nutrient-specific states. To minimize potential downregulation of responding or performance asymmetry induced by the initial extinction tests, all rats were subsequently given a further 5 days of training under general food deprivation prior to the second set of choice tests.

#### Data analysis

As with Experiment 1, the dependent variables from the critical tests were mean lever press rates per minute. Data from the first a second round of choice extinction testing were analyzed using mixed-model ANOVAs followed by simple effects analyses similar to the approach taken in Experiment 1.

## Results and Discussion

### Performance during instrumental training

Instrumental performance during each of the training sessions conducted in Experiment 2 is presented in Figure 12. Responding in Group Whey-high is presented in the left-hand figures and in Whey-low in the right-hand figures. As these graphs show, the Whey-high rats pressed at a higher rate for whey than polycose during the first 10 days of training when pre-fed high carbohydrate meals (Figure 10a) whereas, after a few sessions of training, the rats in Group Whey-low pressed more for polycose than for whey (Figure 10b). After the first round of testing this pattern quickly inverted when the nutrient appetites were switched from protein to carbohydrate hungry, in Group Whey-high, and from carbohydrate to protein-hungry in Group Whey-low (Figures 10c & 10d). Finally, responding during the sessions conducted generally food-deprived were effective in producing higher rates of performance prior to the second round of choice tests and, by the end of the five days of training were roughly similar on the two levers and across the two groups.

**Figure 10.**
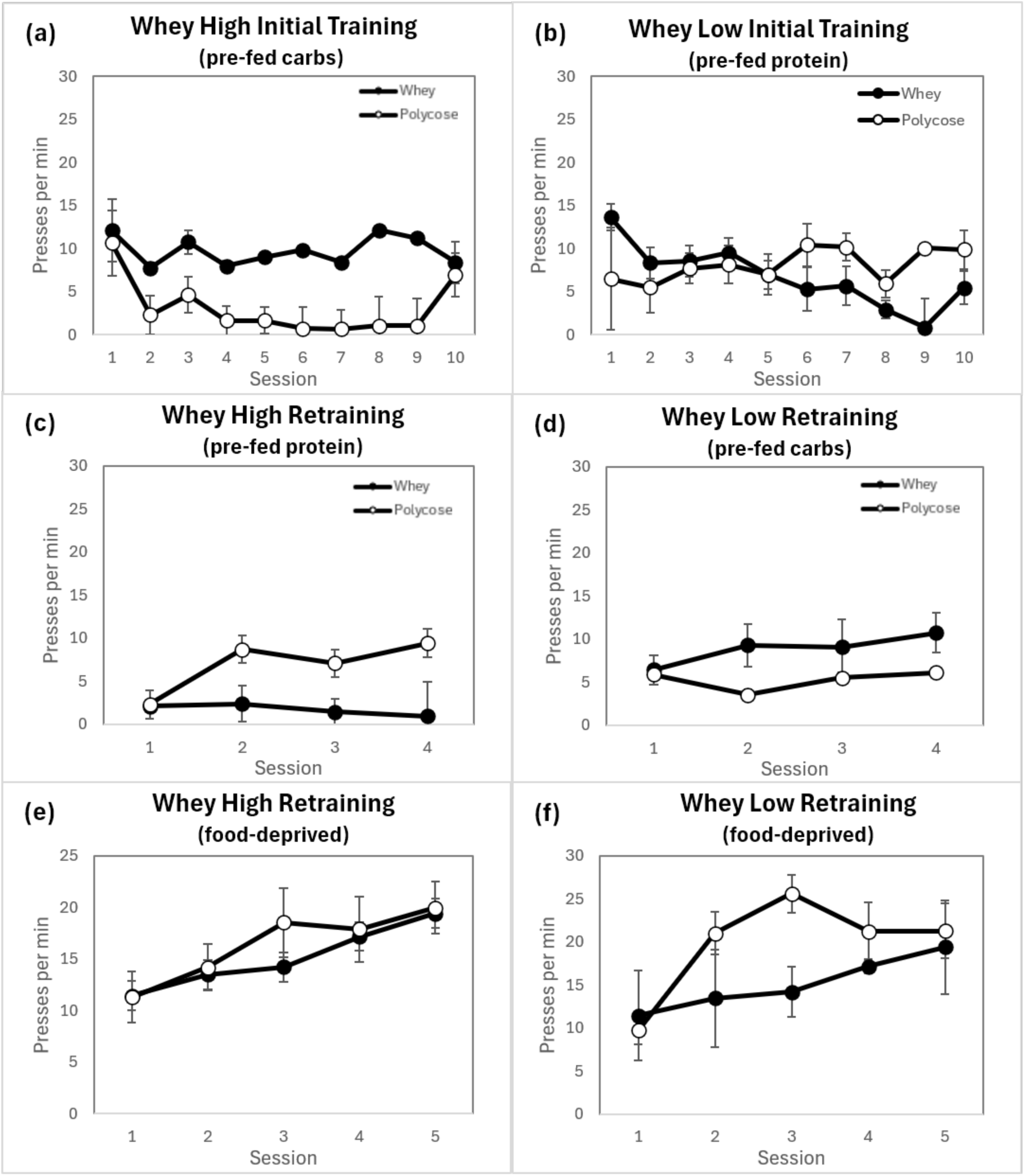
Responses on the whey and polycose levers during the instrumental training and re-training periods for Groups Whey-high and Whey-low. Average lever press rates for each session during initial instrumental training are shown for **(a)** Group Whey-high and **(b)** Group Whey-low under, respectively, protein or carbohydrate appetite prior to the first round of choice tests. Average lever press rates per session are shown for the first retraining period for **(c)** Group Whey-high and **(d)** Group Whey-low under carbohydrate or protein appetite respectively and during the second retraining period for **(e)** Group Whey-high and **(f)** Group Whey-low under general food deprivation prior to the second round of choice extinction tests (Error bars = ±1 SEM).

**Figure 11.**
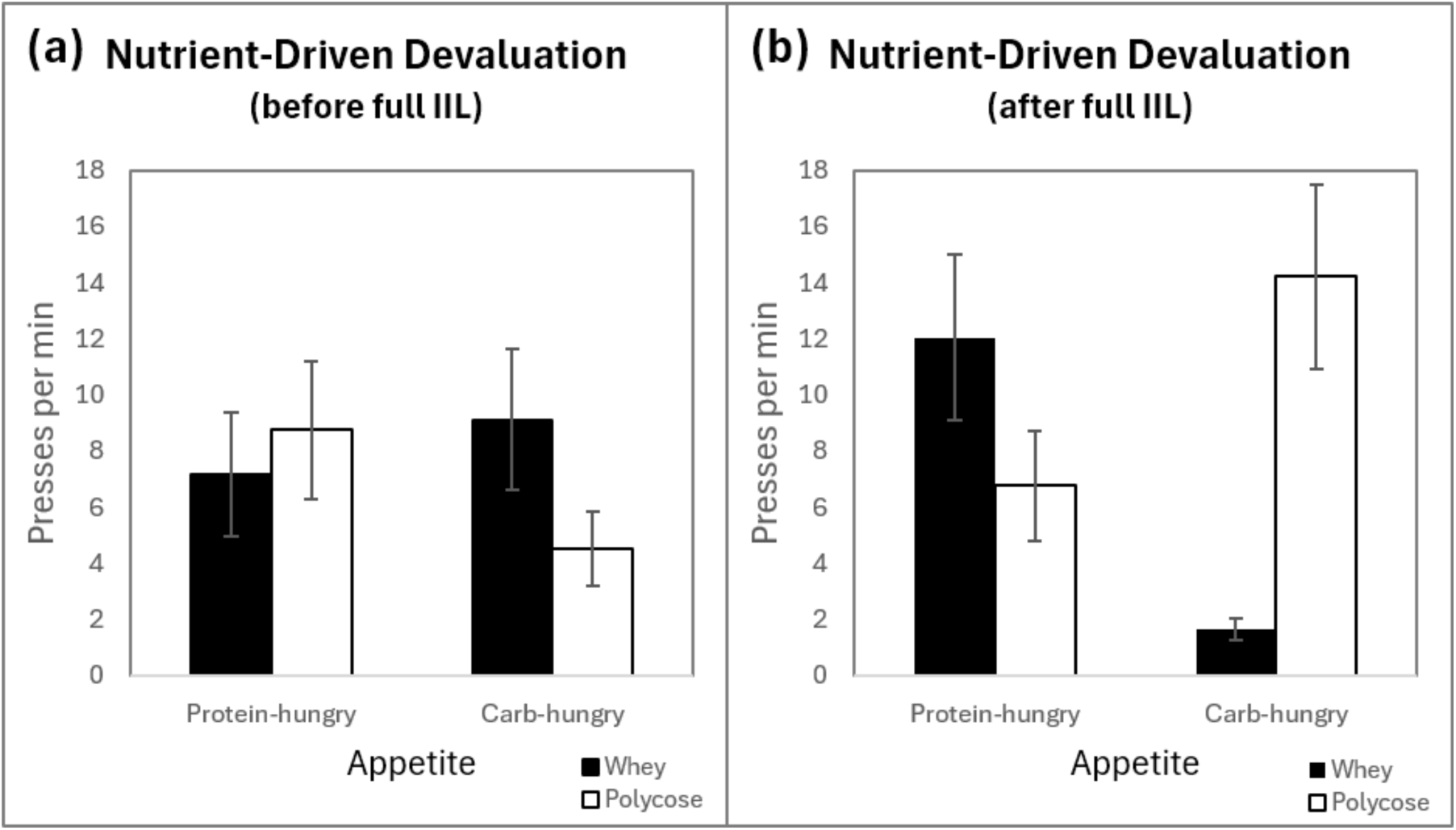
Lever press rates on the whey vs. polycose levers under protein and carbohydrate appetites during extinction choice tests after partial and full incentive learning. Mean lever press rates for the whey protein lever vs. polycose lever across the two appetite conditions **(a)** prior to and **(b)** after full instrumental incentive learning (error bars = ±1 SEM).

**Figure 12.**
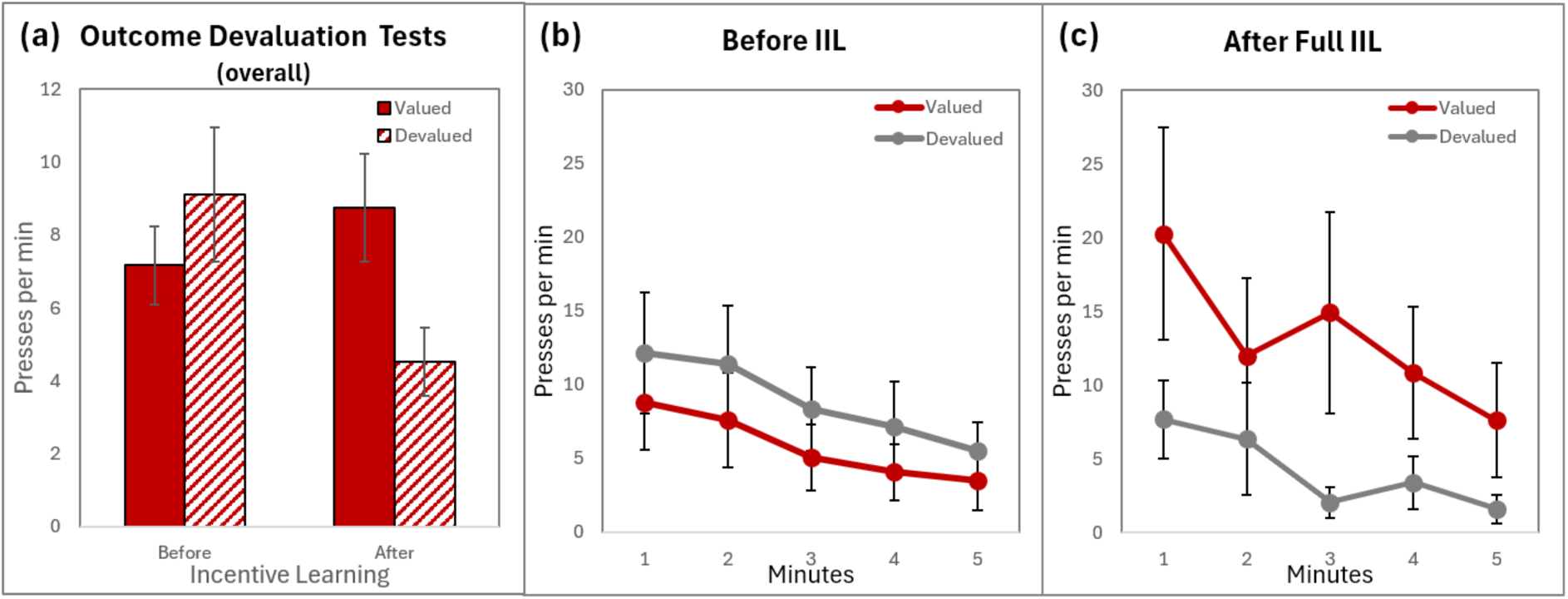
Overall lever press rates averaged across appetites during the extinction choice tests given before and after full incentive learning. **(a)** Mean lever press rates on the valued (congruent with nutritive state at test) and devalued (incongruent with nutritive state at test) levers when the outcome had been devalued after partial and full incentive learning. Performance broken down by minute **(a)** prior to and **(b)** after full instrumental incentive learning (error bars = ±1 SEM).

This description was largely supported by the statistical analysis. Separate two-way ANOVAs were conducted within each group and within each instrumental training phase, with Outcome (whey vs. polycose) and Session (early vs. late) as factors. Prior to the first set of devaluation tests rats in Group Whey-high group pressed more for whey protein (*M*=10.61, *SD =* 7.72) than polycose (*M* = 3.03, *SD =* 2.39), main effect of Outcome: *F*(1,7) = 14.774, *p*=.006, 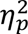 = .679. They then pressed at a lower rate for protein (*M* = 1.60, *SD =* 1.93) than carbohydrate (*M* = 8.40, *SD =* 6.27) when subsequently trained under conditions of protein satiety, *F*(1, 7) = 7.057, *p* = .033, 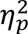 = .502, whereas there was no significant difference in press rates for either outcome when trained under conditions of general food deprivation (no main effect of Outcome: *F*(1, 7)=.003, *p* = .957, 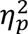 < .001). Similarly, rats in Group Whey-low group pressed on average at a lower rate across sessions for whey (*M* = 3.15, *SD =*3.45) than for polycose (*M* = 8.89, *SD =* 4.90), during initial training; main effect of Outcome: *F*(1, 7) = 9.651, *p* = .017, 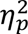 = .580, although during re-training, there was no main effect of Outcome when trained either under protein hunger (*F*(1, 7) = 3.168, *p* = .118, 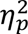 = .312), or general food deprivation (*F*(1, 7) = .076, *p* = .791, 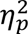 = .011).

#### Responding during the choice extinction tests

The critical choice test data are presented in Figure 11, with performance on each lever (whey or polycose) examined under the different appetite conditions (protein or carb hungry), separated by incentive learning experience at the time of testing (partial or full). There was a clear shift in the pattern of responding across the two sets of tests. During the first set of choice tests, shown in Figure 11a, there was no apparent evidence of control over performance by the nutrient-specific states; the rats did not clearly elevate performance for whey over polycose when protein hungry nor did they elevate responding for polycose over whey when carbohydrate hungry. Indeed, numerically they appeared to show the opposite pattern of responding, something that broadly follows their performance during initial training. In contrast, in the tests conducted after the nutrient appetites were switched from initial training to the re-training period in the two groups, the control by nutrient-specific state was clearer and indeed, appeared to reflect the expected values of the whey and polycose rewards under the nutrient-specific states, Figure 11b. Now rats responded more on the whey lever than the polycose lever when protein hungry and more on the polycose lever than the whey lever when carbohydrate hungry.

Lever pressing per minute during the extinction choice tests was analysed using three-way repeated measures ANOVA with Appetite (protein hungry vs. carbohydrate hungry), Outcome (whey vs. polycose) and Test (first test after partial incentive learning and second test after full incentive learning) as factors. This analysis found no significant main effect of Appetite (*F*(1, 15) = .709, *p* = .426, 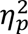 = .045), Outcome (*F*(1, 15) = .232, *p* = .637, 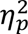 = .015), or Test (*F*(1, 15) = 2.604, *p* = .127, 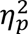 = .148). The Appetite × Test interaction was not significant (*F*(1, 15Te) = .009, *p* = .926, 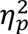 = .001), nor was the Outcome × Test interaction (*F*(1, 15) = 2.373, *p* = .144, 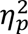 = .137). However, there was both a significant Appetite × Outcome interaction, *F*(1, 15) = 6.658, *p* = .021, 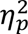 = .307, and, more importantly, a significant three-way interaction between Appetite × Outcome × Test was observed, *F*(1, 15) = 21.898, *p* < .001, 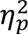 = .593. This crucial three-way interaction supports the claim that that the control of lever press performance by nutritive state depended on both outcome type and whether animals had been exposed to the outcomes under more than one state; i.e., the motivational control of instrumental performance was once again found to depend on incentive learning.

To characterise performance at each stage of training, we compared mean press rates on levers targeted for devaluation to those that were not for each set of choice tests using 2 (Appetite) × 2 (Outcome) repeated-measures ANOVAs. These revealed that, after initial training, there was a significant interaction, *F*(1, 15) = 4.756, *p* = .046, 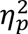 = .241. However, simple effects failed to find any significant difference in either pressing for protein (*F*(1, 15) = 2.329, *p* = .148, 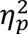 = .134) or carbohydrate under different appetite conditions (*F*(1, 15) = 2.3, *p* = .150, 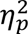 = .133).

By contrast, following full incentive learning, a robust Appetite × Outcome interaction was detected *F*(1, 15) = 20.693, *p* < .001, 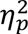 = .580, with simple effects confirming that these post-full IIL results were characterised by distinct behavioural shifts within this second test phase. Specifically, when protein hungry, rats pressed the whey protein lever more times per minute (*M* =12.00, *SD* = 11.76) than when carbohydrate hungry (*M* = 1.65, *SD* = 1.54) *F*(1, 15) = 12.656, *p* = .003, 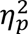 = .458. Conversely, carbohydrate hunger only marginally affected presses on the lever associated with polycose (*M* = 14.21, *SD* = 13.12) compared to the protein appetite (*M* = 6.75, *SD* = 1.94) (*F*(1, 15) =3.520, *p* = .082, 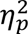 = .178). Finally, when looking at within-test lever bias during these tests after full incentive learning, the protein appetite failed to significantly bias performance on the levers within a test (*F*(1, 15) = 3.444, *p* = .083, 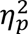 = .187), whereas the carbohydrate appetite produced a significant bias in performance such that the rate of lever pressing was greater for the polycose than the whey lever, *F*(1, 15) = 14.020, *p* = .002, 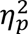 = .483.

Finally, to compare the two sets of tests more directly, we collapsed data across the relative value of the outcomes earned by the actions under the two nutrient appetites on test to express performance on each lever as either valued or devalued for the tests conducted before and after full incentive learning (see Figure 12). A repeated measures ANOVA was conducted to compare factors of Value (valued vs. devalued) across the two Tests (before and after full incentive learning). There was no main effect of Test suggesting that the intervening period did not change how much rats pressed overall during these extinction choice tests (*F*(1, 15) = .2553, *p* = .132, 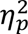 = .154), however there was a main effect of Value, *F*(1, 15) = 6.402, *p* = .024, 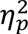 = .314, and, importantly, a significant Value x Test interaction, *F* (1, 15) = 22.199, *p* <.001, *ηp²* = .613. Simple effects analyses revealed that, after partial incentive learning, rats pressed the lever associated with reward targeted for devaluation at a *higher* rate than the valued lever, *F*(1, 15) = 4.756, *p* = .046, 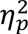 = .241. However, after full incentive learning, the rats’ performance revealed significant nutrient-specific control of value, pressing the lever associated in training with the valued outcome more than the action associated with the devalued outcome, *F*(1, 15) = 20.693, *p* < .001, 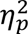 = .580.

Generally, therefore, the results of Experiment 2 confirmed that nutrient-specific appetites control instrumental performance both by determining the incentive value of specific outcomes and via the conditional control of value, the latter driven by the differential retrieval of outcome values based on appetite-specific cues.

## General Discussion

Our results generally replicate Roy et al.’s (2026) finding that rats moderate their instrumental efforts based on the specific nutrients a reward contains. Pre-feeding rats that were otherwise food-deprived with a meal high in protein (but lacking carbohydrate) selectively reduced press rates on a lever they had learned to cause delivery of whey protein, while pre-feeding them a high carbohydrate meal (lacking protein) selectively influenced their rates of pressing a lever for a high carbohydrate reward. However, for this nutrient-driven devaluation effect to become manifest, we found that rats require experience with these rewards under different degrees of appetite: when we restricted the range of experiences rats had with outcomes across appetite conditions, the nutrient-driven devaluation effect was either distorted or prevented entirely. For instance, when whey had only ever been encountered under conditions where it would have been highly rewarding, rats continued to press for it at rates commensurate with it having a relatively high value *regardless* of whether they were acting under a state of protein appetite or satiety. This dependency on prior experience indicates that mere exposure to the rewards is not sufficient for motivational control of goal-directed action by nutrient specific appetites.

These findings are difficult to reconcile with a structural occasion-setting account (Fraser & Holland, 2019) in which appetites selectively modulate response performance directly. By isolating incentive learning to the home cage in Experiment 1, we dissociated the acquisition of incentive value from the instrumental context, suggesting instead that appetites modulate the expected value of outcome representations, which animals then use to guide instrumental choice. This account of our findings leads us to suggest that the learning underpinning the nutrient-driven devaluation effect is best understood as an example of instrumental incentive learning (Dickinson & Balleine, 1994; Balleine, 2000) with patterns of behaviour we observed during the extinction choice tests echoing the standard incentive learning asymmetry for hunger vs. satiety (e.g., Balleine, 1992; Dickinson & Balleine, 1994).

### Implications

An implication of our findings is that the results reported by Roy et al (2026) likely also depended on the sort of experience given to subjects to familiarize them with rewards under different motivational conditions. This interpretation complements the view that performance in those tests reflected rats’ expectations about the current value of the outcomes given their nutrient-specific appetites and was therefore goal-directed rather than stimulus-driven. In this respect, nutrient-driven devaluation appears to operate similarly to standard outcome revaluation procedures in instrumental conditioning (e.g., Adams & Dickinson, 1981; Dickinson & Balleine, 1994).

Mapping these cognitive functions onto the brain introduces a distinct set of neurobiological questions. If our results represent a nutrient-specific instance of instrumental incentive learning more generally, it may depend on neural systems already implicated in incentive learning and goal-directed action, including the insular cortex and corticostriatal valuation networks (e.g., Balleine & Dickinson, 2000). Recent evidence that protein deprivation alters activity within mesolimbic reward systems, including the nucleus accumbens and ventral tegmental area (Chiacchierini et al., 2021; Tomé et al., 2019), further suggests that nutrient-specific appetites may modulate the neural representation of outcome values. Future work will be required to characterise how nutrient-specific motivational states interact with these systems to calibrate goal-directed action and choice.

Beyond its neural underpinnings, this incentive learning framework offers a different angle for understanding how individuals adjust (or fail to adjust) as nutritional requirements shift over time. If nutrient-specific appetites guide goal-directed behaviour through incentive learning, then effective dietary adaptation to nutritional challenges related to aging, illness, physical activity, shifts in body composition, and so on, may depend partly on an individual’s ability to update the expected value of foods as their nutritional needs change. Failures or distortions in this updating process could conceivably contribute to persistent behavioural patterns that are no longer adaptive, and nutrient-specific incentive learning could be a crucial mechanism for helping individuals align their behavioural responses to underlying physiological requirements.

This vulnerability to updating failures becomes particularly acute when evaluating modern obesogenic environments. For instance, the ‘protein leverage hypothesis’ proposes that restriction to protein-dilute diets promotes excess energy intake (Simpson & Raubenheimer, 2005, 2012). One challenge for this account is explaining why humans living in modern food environments often fail to consistently select foods that efficiently satisfy protein requirements despite widespread dietary availability. One possibility is that modern diets disrupt or obscure the normal incentive learning processes through which nutrient-specific appetites calibrate goal-directed food choice. Understanding how nutrient-specific incentive learning develops, generalizes, and adapts across changing dietary environments may therefore prove useful for understanding both obesity and other disorders linked to dysregulated feeding behaviour, illuminating why individuals in modern food landscapes often forage as if choices are scarce despite an unprecedented abundance of dietary options.

### Future directions

While the foregoing helps to explain how experience informs appetite-related actions, some methodological constraints would benefit from further clarification by future research. Firstly, the current studies focused specifically on protein and carbohydrate appetites. Determining whether similar incentive learning processes govern responses to dietary fats, micronutrients, or more specific nutrient imbalances will be important for establishing the broader generality of these effects. Many taxa can detect and respond to highly specific nutritional deficits (Simpson & Raubenheimer, 2012), including imbalances in essential amino acids (Gietzen, 2022), raising the possibility that incentive learning processes may operate across multiple levels of nutritional precision. Understanding how experience shapes the calibration, generalization, and accuracy of nutrient-specific incentive learning may therefore prove important for explaining both adaptive and maladaptive patterns of food-directed behaviour.

Secondly, the present findings were limited to Long Evans rats, and replication across other species with differing foraging ecologies will be necessary to determine the phylogenetic scope of our findings. At the time of writing, there is a lack of direct demonstration of instrumental incentive learning in humans. Recent technological innovations that allow the macronutrient profile of foods and drinks to be manipulated largely independently of superficial sensory characteristics may make it viable to use nutrient-specific appetites to explore instrumental incentive learning processes in humans. It should be relatively straightforward to induce specific nutrient appetites in human participants and manipulate their experiences with sufficiently novel rewards similar in nutritional composition to those used in the rat experiments discussed here, before testing them in choice situations to see how they allocate their efforts between actions earning these rewards.

Thirdly, while our findings point strongly toward an incentive learning account of nutrient-driven devaluation rather than a structural occasion-setting mechanism, a separation of these processes would be strengthened by further tests. Another way to achieve this would be to directly manipulate the action–outcome contingency (Balleine & Dickinson, 1998). Future studies could explore this along the following lines. Suppose animals are initially trained to press one lever for whey protein and another for polycose. If another source of protein were then delivered (e.g., pellets high in beef protein) in a manner that was not contingent on lever presses, we should expect this to selectively degrade pressing on the whey lever more than the polycose lever: if the animal is pressing for whey because they are hungry for protein, then getting protein from another source makes pressing for whey less necessary. Crucially, if such nutritional factors affect behaviour in this way, we may further predict that, like the nutrient-driven devaluation effect, animals must have had instrumental incentive learning about the outcomes; i.e., they must have learned through experience that beef pellets are relevant to protein appetite for free deliveries of beef pellets to attenuate actions earning whey protein.

## Summary and Conclusions

Overall, these experiments show that nutrient-specific appetites guide goal-directed action only after animals have had sufficient experience of the value of those outcomes under different nutrient states. When experience with protein and carbohydrate rewards was restricted across appetite conditions, rats failed to adjust their responding appropriately to current nutritional need; when such experience was available, their choice behaviour tracked the current motivational significance of each outcome. This pattern suggests that nutrient-driven devaluation depends on learning the expected value of outcomes across states of appetite rather than on direct stimulus control by the appetite itself. More broadly, the results support an instrumental incentive learning account of nutrient-specific motivation and raise the possibility that adaptive food-directed behaviour depends on updating outcome value in light of changing physiological needs. More research is needed to determine how widely this mechanism generalizes across nutrients, species, and human dietary contexts, and what factors affect its accuracy. Nevertheless, the current results are important in significantly expanding our understanding of the scope of incentive learning effects and its general importance in the motivational control of goal-directed action.

## ACKNOWLEDGEMENTS

This research was supported by an award from the Australian Government Research Training Program to Douglas Roy and Australian Research Council grants DP200103401 to Bernard Balleine and DP240103246 to Bernard Balleine and Thomas Burton. The authors declare no conflicts of interest.

